# Post-Transcriptional Methylation of Mitochondrial-tRNA Differentially Contributes to Mitochondrial Pathology

**DOI:** 10.1101/2023.12.09.569632

**Authors:** Sunita Maharjan, Howard Gamper, Yuka Yamaki, Robert Y. Henley, Nan-Sheng Li, Takeo Suzuki, Tsutomu Suzuki, Joseph A. Piccirilli, Meni Wanunu, Erin Seifert, Douglas C. Wallace, Ya-Ming Hou

## Abstract

Human mitochondrial tRNAs (mt-tRNAs), critical for mitochondrial biogenesis, are frequently associated with pathogenic mutations. These mt-tRNAs have unusual sequence motifs and require post-transcriptional modifications to stabilize their fragile structures. However, whether a modification that stabilizes a wild-type (WT) mt-tRNA structure would also stabilize its pathogenic variants is unknown. Here we show that the *N*^1^-methylation of guanosine at position 9 (m^1^G9) of mt-Leu(UAA), while stabilizing the WT tRNA, has an opposite and destabilizing effect on variants associated with MELAS (mitochondrial myopathy, encephalopathy, lactic acidosis, and stroke-like episodes). This differential effect is further demonstrated by the observation that demethylation of m^1^G9, while damaging to the WT tRNA, is beneficial to the major pathogenic variant, improving its structure and activity. These results have new therapeutic implications, suggesting that the *N*^1^-methylation of mt-tRNAs at position 9 is a determinant of pathogenicity and that controlling the methylation level is an important modulator of mt-tRNA-associated diseases.

## Introduction

Mitochondria are essential eukaryotic organelles that produce most of the cellular ATP molecules through the process of oxidative phosphorylation. This mitochondrial energy production requires a balanced mechanism that integrates the multicopy mt-DNA genome with the nuclear genome. Specifically, while the mt-DNA expresses a small number of mt-proteins, the vast majority of mt-proteins are expressed from the nuclear genome, synthesized in the cytosol, and imported into the organelles. In humans, due to the high demand for aerobic activity and developmental complexity, the mt-DNA has accumulated much higher levels of mutations as compared to the nuclear genome.^1–4^ While mt-DNA mutations induce stress and dysfunction, which activates cellular surveillance mechanisms, chronic perturbations lead to pathologies.^5^ Importantly, most of the disease-causing mt-DNA mutations are mapped to mt-tRNAs (MITOMAP, www.mitomap.org)^6,7^ and are associated with devastating neurological disorders and cardiovascular myopathies. The strong association of mt-tRNAs with mitochondrial pathologies underscores the need for a better understanding of these tRNAs to improve the well-being of the public.

The human mt-DNA encodes genes for 22 mt-tRNAs, 2 mt-rRNAs, and 13 mt-proteins.^8,9^ All 22 mt-tRNAs are used in the mt-ribosome machinery, which is assembled from the 2 mt-rRNAs and nuclear-encoded mt-proteins.^8,10^ Each of the 13 mt-proteins is a core component of the mitochondrial electron transport chain that produces ATP,^11^ emphasizing the importance of each mt-tRNA in its fully functional state to contribute to mitochondrial protein synthesis and information processing. Notably, mt-tRNAs are transcribed with inherent structural fragility, having low sequence complexity, weak base pairs and mismatches in stem regions, small loop sizes, loss of conserved tertiary interactions from the canonical tRNA structure, and even deletions of entire domains, contributing to their generally low thermal stability.^8,12,13^ This structural fragility is mitigated in each mt-tRNA by acquiring a defined set of post-transcriptional modifications,^14^ all of which are catalyzed by nuclear-encoded enzymes. These enzymes are frequently associated with human pathologies as well,^15^ constituting another layer of mitochondrial biology at the nexus of the mitochondrial and the nuclear genomes.

The most prominent post-transcriptional modification in mt-tRNAs is the *N*^1^-methylation to the purine R (A/G) nucleotide at position 9, which is conserved in 19 of the 22 mt-tRNAs (5 as m^1^G9 and 14 as m^1^A9),^16^ whereas the remaining 3 mt-tRNAs each contains a pyrimidine at position 9. The m^1^G9/m^1^A9 methylation is catalyzed by a nuclear-encoded ternary enzyme complex that has dual-specificity for both purine nucleotides. This ternary complex consists of MRPP1-2-3 (mitochondrial RNase P proteins 1, 2, and 3), where MRPP1 (*a.k.a*,, TRMT10C) is the methyl transferase, MRPP2 (SDR5C1) is the short-chain oxidoreductase/hydroxysteroid 17β-dehydrogenase 10, and MRPP3 (PRORP) is the 5’-processing enzyme of all mt-tRNAs.^17–19^ As the 5’-processing catalyzed by the MRPP1-2-3 complex is the initiation step of maturation of all mt-tRNAs,^19,20^ it is widely believed that the associated m^1^G9/m^1^A9 methylation is installed at the time of 5’-processing as the first modification that folds each mt-tRNA into a structure suitable for subsequent modifications.^21^ Indeed, while the transcript of human mt-Lys(UUU) (UUU, the anticodon), lacking any modification, adopts an extended and non-functional stem-loop structure, introduction of the m^1^A9 methylation to the transcript converts it to a structure closely similar to the native-state.^22–24^ Thus, of all natural modifications present in human mt-Lys(UUU), the single m^1^A9 is sufficient to fold a non-functional tRNA into a functional state that sets the stage for additional modifications. Notably, this finding was based on the wild-type (WT) sequence of mt-Lys(UUU), but not a pathogenic variant. We do not yet know whether the *N*^1^-methylation of purine 9 would have the same, or a different, effect on a pathogenic variant. Given that many pathogenic variants are unstable,^25,26^ due to mutations that further weaken an already fragile structure, they might benefit from the *N*^1^-methylation of purine 9 to strengthen the structure.

Here we address this question directly, using human mt-Leu(UAA) as a model (Figure 1A), which has m^1^G9 (Figure 1B) as shown in an LC-MS analysis.^16^ This mt-tRNA was previously denoted as mt-tRNA^Leu^(UUR), indicating its reading of the leucine codons UUR (R: A/G) during protein synthesis in mitochondria. We chose mt-Leu(UAA) as a different sequence framework than mt-Lys(UUU) to more broadly test the importance of the *N*^1^-methylation of purine 9. Notably, mt-Leu(UAA) has been examined by structural probing and aminoacylation experiments,^27,28^ and is one of the few human mt-tRNAs possessing all of the conserved features for the canonical tRNA structure (Figure 1A).^29^ A previous study showed that the transcript of WT mt-Leu(UAA) has a weakly stable structure, sampling between a folding intermediate and the native-state, but that the addition of all natural post-transcriptional modifications to the transcript produces a more constrained native structure that does not sample the intermediate conformation.^30^ While this work emphasizes the importance of tRNA modifications, it does not point to a specific modification as the major driver of folding. Importantly, human mt-Leu(UAA) is most closely associated with MELAS.^31^ Approximately 80% of patients with MELAS have the mt-DNA A3243G substitution that is mapped to mt-Leu(UAA),^32,33^ representing the most common single-point pathogenic mutation in human populations.^34^ Another 10% of patients with MELAS have the T3271C substitution that is also mapped to mt-Leu(UAA) (Figure 1A) as the second most common mutation associated with the disease.^35^ The A3243G substitution corresponds to the A14G mutation in mt-Leu(UAA), which disrupts the 8-14 reverse-Hoogsteen base pair that connects the D loop with the tight turn from the acceptor stem to the D stem in the canonical tRNA structure. Separately, the T3271C substitution corresponds to the U40C mutation in mt-Leu(UAA), which disrupts the 30-40 base pair of the anticodon stem. The two mutations thus represent two distinct structural perturbations of the same mt-tRNA. However, despite the close association of these mutations with MELAS, the molecular basis of the pathology remains unclear. While both the A3243G and T3271C variants of mt-Leu(UAA) share common features – deficiency in the level of tRNA stability, aminoacylation, and the 5-taurine-methyl modification to the wobble nucleotide U34 (ρm^5^U34)^36^ (Figure 1C), the molecular basis of these deficiencies is unknown.

**Figure 1.**
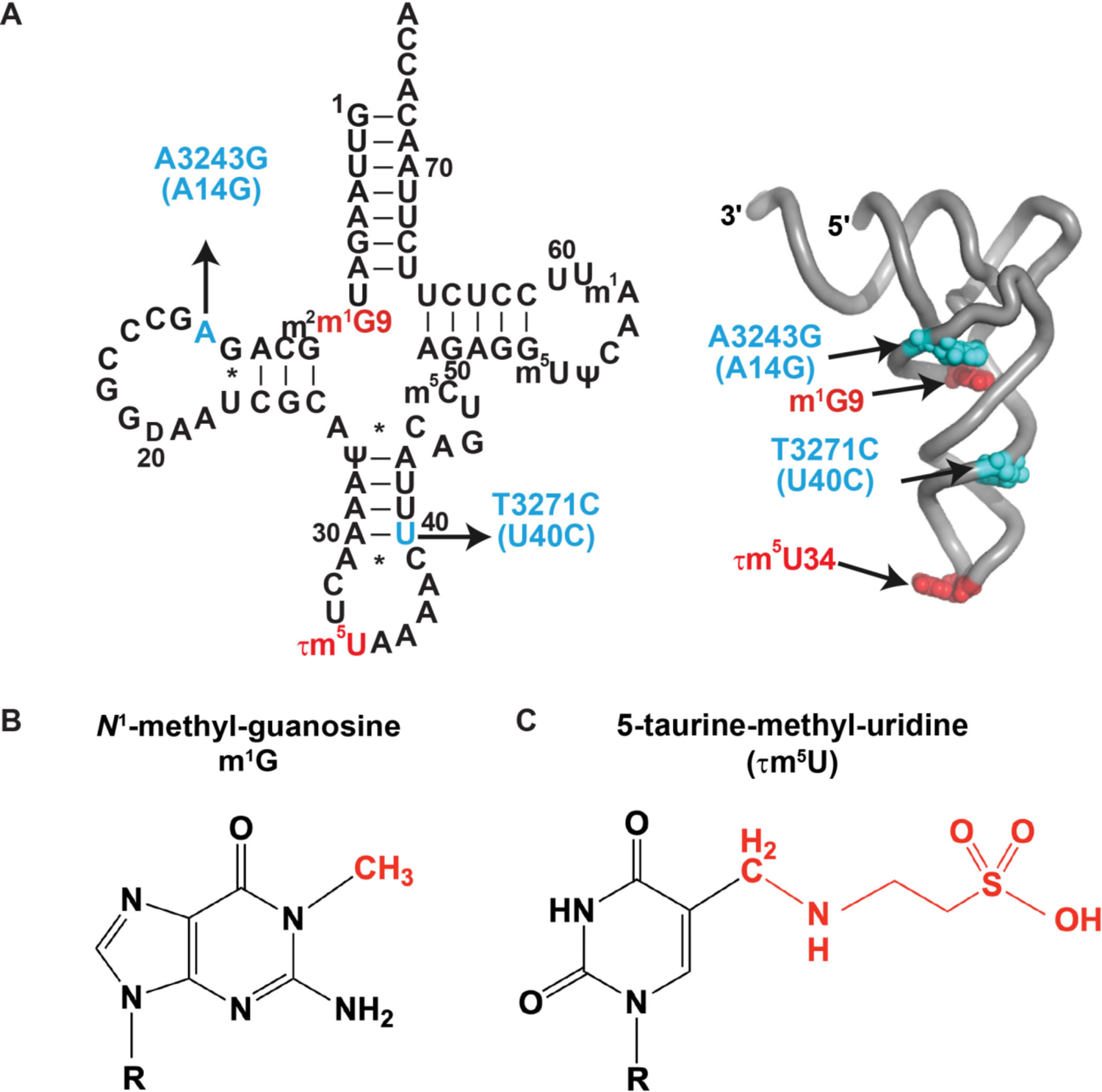
Sequence and post-transcriptional modifications of human mt-Leu(UAA) (A) Sequence and cloverleaf structure of human mt-Leu(UAA) showing the A3243G and T3271C substitutions in blue and the m^1^G9 and 1m^5^U34 modifications in red, both mapped to the canonical L-shaped tRNA structure. (B) Chemical structure of m^1^G, where the *N^1^*-methyl group is in red. (C) Chemical structure of 1m^5^U, where the taurine-*C^5^*-methyl group is in red.

We set out to functionally separate the m^1^G9 methylation in human mt-Leu(UAA) from all other natural modifications to elucidate its role in the disease mechanism of MELAS. Unexpectedly, we show that, while m^1^G9 indeed stabilizes the WT mt-Leu(UAA) structure, it has an opposite effect on both the A3243G and T3271C variants, trapping each in an aberrant structure. Extensive analysis of the A3243G variant reveals that the m^1^G9-induced aberrant structure is unstable and is sensitive to degradation. Conversely, we show that demethylation of m^1^G9 is beneficial to the A3243G variant in mitochondria, whereas it is damaging to the WT tRNA. Further, demethylation of m^1^G9 in cells partially restores mitochondrial respiration of the variant relative to the WT. These results produce a new conceptual framework that emphasizes the ability of m^1^G9 to differentially control and regulate mitochondrial health and pathology. This framework has a novel therapeutic implication in that, instead of enhancing m^1^G9 in mt-Leu(UAA) as a remedy for MELAS, it suggests the opposite. Given the broad conservation of m^1^G9/m^1^A9 in human mt-tRNAs, and the high-stoichiometry of the methylation even in pathogenic variants of mt-tRNAs, this framework is likely generalizable to other mt-tRNA-associated diseases.

## Results

### The m^1^G9 methylation in mt-Leu(UAA) is the major driver to reach the native structure

We addressed whether m^1^G9 is the major driver that determines the folding of human mt-Leu(UAA) to the native structure. To determine folding, we measured the electrokinetic translocation of the tRNA through solid-state nanopores using our previously demonstrated label-free single-molecule assay.^37^ This measurement distinguishes different RNA structures by the associated pore translocation times and ion-current signals.^38^ In the nanopore device (Figure 2A, top), as a voltage is applied across a small synthetic pore (∼3 nm diameter), a tRNA molecule is electrophoretically driven to unfold and to translocate through an electrolyte filled pore, displacing some of the ions as it passes through, thus lowering the open pore current in an event defined by two parameters.^38^ The parameter “current blockade” is the difference between the open pore current and the measured current amplitude, while the parameter “dwell time” is the duration of the molecule within the pore. These two parameters provide structural information on the tRNA molecule passing through the pore, such as the shape, size, and the conformation dynamics in both folded and unfolded states.^39,40^ For example, a properly folded tRNA typically resides in the pore longer until it eventually unfolds and passes through the pore with a higher current blockade, whereas an improperly folded tRNA passes through the pore faster with a lower current blockade.^39,40^

**Figure 2.**
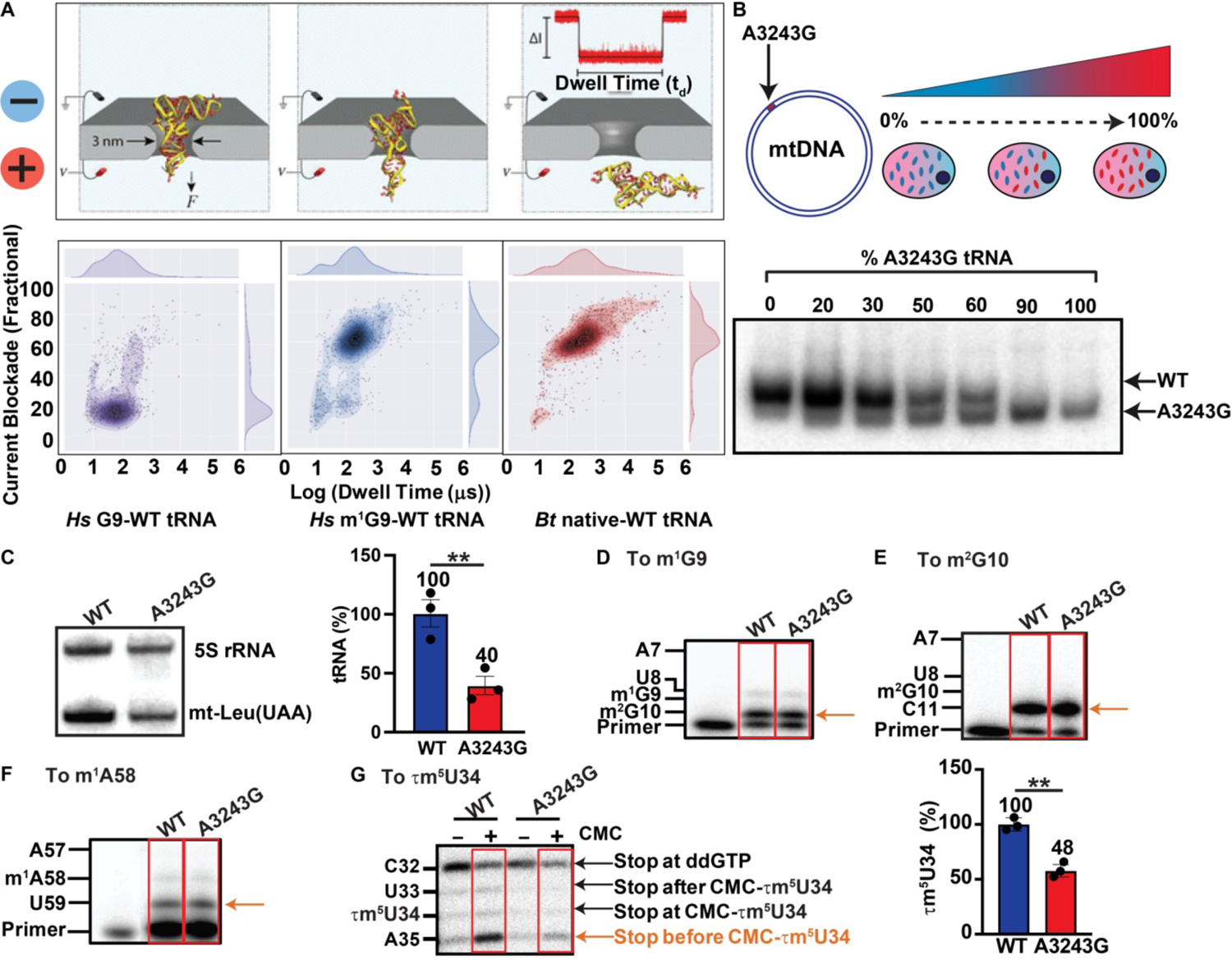
The A3243G variant has an altered tRNA structure, despite having m^1^G9 (A) Top: Electrophoretic unfolding of a tRNA through a solid-state nanopore. Schematics show steps required for tRNA passage through a 3-nm diameter nanopore, along with an example of a translocation event. Bottom: Heat-map scatter plots of current blockade as a function of dwell time (log_10_) of the WT transcript of human mt-Leu(UAA) in the G9-state (left), the WT transcript of human mt-Leu(UAA) in the m^1^G9-state (middle), and the *Bos taurus* liver mt-Leu(UAA) in the native-state with the full-complement of all natural modifications (right). (B) Top: A series of A3243G cybrids with an increasing frequency of heteroplasmy. Bottom: A native 12% PAGE separation of the WT from the A3243G variant of human mt-Leu(UAA) in the heteroplasmy series as probed by Northern blot analysis of the tRNA. (C) A Northern blot analysis of the steady-state level of human mt-Leu(UAA) relative to the 5S rRNA in the WT and A3243G cybrids. Data in the bar graph represent the average ± SD, \*\**p*<0.01 (n = 3). (D) A representative primer extension analysis of the m^1^G9 level on total RNA isolated from WT and A3243G cybrids. The region of interest is shown by a red box and the band due to termination of primer extension is indicated by an orange arrow. (E) A representative primer extension analysis of the level of m^2^G10 on total RNA isolated from WT and A3243G cybrids. The red box and the orange arrow are as in (D). (F) A representative primer extension analysis of the level of m^1^A58 on total RNA isolated from WT and A3243G cybrids. The red box and the orange arrow are as in (D). (G) A representative analysis of the level of 1m^5^U34 on total RNA isolated from WT and A3243G cybrids by CMC-assisted primer extension. Total RNA was treated with CMC (+) or buffer (−). The level of 1m^5^U34 was calculated as the fraction of the band at A35 in the sum of the bands at A35, 1m^5^U34, U33 and C32. Data in the bar graph are average ± SD, \*\**p*<0.01 (n = 3). SD is standard deviation and p value was obtained from Student’s t test.

To test if m^1^G9 is the driver in folding of human mt-Leu(UAA), and to separate it from all other natural modifications in the tRNA, we prepared mt-Leu(UAA) with the WT sequence in two states, the transcript G9-state lacking any modifications, and the m^1^G9-state containing the single m^1^G9 modification in the transcript-state. Each state was allowed to fold in the presence of Mg^2+^ and to then pass through nanopores. Measurement of current blockade as a function of dwell time (in Log_10_ scale) showed that the transcript G9-state had a low fraction of current blockade (20%) and a short dwell time (log_10_2 μs) (Figure 2A, bottom, left), but that the m^1^G9-state had a higher fraction of current blockade (70%) and a longer dwell time (log_10_2.4 μs) (Figure 2A, bottom, middle). The increase in the current blockade indicates that the single m^1^G9 methylation transformed the transcript G9-state to a more stable structure that required higher energetics of unfolding during passage through the pore. Notably, the translocation profile of the m^1^G9-state was quantitatively similar to that of the native mt-Leu(UAA) isolated from bovine liver (Figure 2A, bottom, right), possessing all natural modifications.^41^ This similarity is notable, due to the similar sequence and modification profile between bovine and human mt-Leu(UAA) in the native-state (Figure S1), indicating that the single m^1^G9 is the major driver that folds human mt-Leu(UAA) into a structure similar to the native structure of the bovine tRNA counterpart.

### A different structure of the A3243G variant, despite having m^1^G9

We used a cybrid cell model to compare the WT and the A3243G variant of MELAS. Cybrids are created by intercellular transfer of the mt-DNA of a patient with a mitochondrial mutation into the mt-DNA-lacking human *r*^0^ cells.^42,43^ Such cybrids have been powerful tools to dissect mitochondrial disorders, including those caused by the A3243G mutation^44,45^ and by others.^46,47^ A unique strength of cybrids is that they provide direct evidence for the association of a specific mt-DNA mutation with the disease pathology,^48^ excluding the role of the nuclear genome in the biogenesis of mitochondria. We previously used the cybrid technique, starting with a patient cell line heteroplasmic of the A3243G mutation, and isolated a series of clones, ranging in the frequency of the mutation (0, 20, 30, 50, 60, 90, and 100%) (Figure 2B).^49^ Using this series of the A3243G variant, where all cybrids were isogenic for the nuclear and mt-DNA genomes, we showed that the progressive increase in the mutation heteroplasmy is correlated with abrupt changes of transcriptional profiling.^49^ Here we used this series to systematically monitor the structural folding of the A3243G variant relative to the WT tRNA. Analysis of total RNA isolated from each cybrid line, followed by Northern blot analysis of mt-Leu(UAA) in a non-denaturing gel, showed that the WT and A3243G variant migrated differently (Figure 2B), indicating a distinct structure of each. Importantly, the two structures distributed relative to each other in a way consistent with the mutation frequency of each cybrid line (Figure 2B), enabling assignment of the variant to the faster migrating species in the gel. In experiments below, unless otherwise noted, we compared between the WT cybrids (0% mutation) and the A3243G cybrids (100% mutation) of this heteroplasmic series.

We provided additional evidence for the variant tRNA adopting a different structure than the WT by showing that it had reduced stability in mitochondria. In a Northern blot analysis of the same amount of total RNA isolated from the WT and A3243G cybrids, we observed that the abundance of the variant tRNA was 40% of the WT (Figure 2C), indicating that it has a less stable structure that was sensitized to degradation. The loss of stability of the A3243G variant tRNA was also reported previously by the same Northern blot analysis.^36,50^

Importantly, while the variant tRNA adopts a different structure than the WT, it has nonetheless retained m^1^G9 in stoichiometry. Due to the limited quantities of mt-Leu(UAA) that we isolated from cybrids, LC-MS measurement of the methylation was not practical. Instead, we probed the methylation status using a primer-extension assay, where the methyl group of m^1^G9 occurring in the W-C face of the guanosine (Figure 1B) would interfere with primer extension by a reverse transcriptase (RT). The nature of the interference varied with the chemical environment of the modification and the processivity of the RT that catalyzed the primer extension. While some RTs would read through the modification, others would stop, generating an extension product that terminated one nucleotide before the modification. We designed the primer extension assay by placing the 3’-end of the primer ∼2 nucleotides before the modification (Figure S2, Table S1), and by including the ddNTP that would terminate the extension at the nucleotide after the modification. We then calculated the fraction of the extension product that terminated one-nucleotide before the modification relative to the sum of all extension products.

We used the presence of m^1^G9 in the WT mt-Leu(UAA) as the guide to screen for RTs that would terminate primer extension in response to the modification. Analysis of total RNA isolated from WT cybrids, we found that the Luna RT enzyme was the only one that extended the 3’-end of a primer from position C11 to m^2^G10 in a reaction containing dCTP and ddATP (Figure 2D, Figure S2A). The RT stop at m^2^G10 indicated that primer extension incorporated a nucleotide opposite m^2^G10 but that it was terminated by the presence of m^1^G9. Analysis of the termination product relative to the readthrough to U8 showed ∼91% of m^1^G9 in samples isolated from WT cybrids (Figure 2D), consistent with an LC-MS analysis.^16^ To confirm that the termination was indeed the readout for m^1^G9, and not for m^2^G10, we performed a primer extension analysis with a series of mt-Leu(UAA) in the transcript-state, lacking m^2^G10 but containing m^1^G9 at 0, 25, 50, and 100% (Figure S2B). A linear correlation between the prepared level of m^1^G9 and the measured level of the RT stop was observed (Figure S2C), indicating that the assay was quantitative with m*^1^*G9. In this titration analysis, lacking m^2^G10, we detected m^1^G9 at the full stoichiometry when the prepared sample was homogeneous with the methylation, which was virtually identical to the level measured in the presence of m^2^G10 in WT cybrids (91%) (Figure 2D). Thus, the readout was independent of m^2^G10 but was responsive to the level of m^1^G9. Applying this assay to total RNA of A3243G cybrids, we observed an RT stop at a level similar to that of WT cybrids (Figure 2D), indicating that the variant has retained a similar stoichiometry of m^1^G9, consistent with previous results.^36^

While the variant contained m^1^G9, we tested whether it had changed other modifications in the tRNA due to its different structure than the WT. Notably, each tRNA is modified in a defined order by modification enzymes, some acting early during the precursor state, while others acting later after processing and during tRNA folding. The different structure of the variant relative to the WT could alter its modification pathway and prevent some modification enzymes from acting. We focused on modifications that are present in the WT tRNA and quantified a subset of these in the variant using the primer extension assay. At m^2^G10, immediately on the 3’-side of m^1^G9, primer extension analysis with SuperScript III (SSIII) identified a similar level of modification in the variant as in the WT (Figure 2E, Figure S2D). At m^1^A58, adjacent to m^1^G9 in the canonical tRNA tertiary structure, primer extension analysis with SSIII also identified a similar level of modification in the variant as in the WT (Figure 2F, Figure S2E). This is unlike the mitochondrial disease MERRF (myoclonus epilepsy with ragged-red fibers), where the A8344G mutation (mapped to A55G) in mt-Lys(UUU) causes loss of the m^1^A58 methylation.^51^

The 1m^5^U34 modification was of interest, as it permits efficient decoding of mt-Leu(UAA) of not only the UUA codon but also the UUG codon for Leu.^52–57^ Indeed, the 1m^5^U34 level is closely correlated with clinical presentation of MELAS, where patients with pathogenic variants deficient of 1m^5^U34 (e.g., A3243G, T3271C) present severe symptoms, while those retaining 1m^5^U34 show few symptoms.^58^ Also, restoration of the 1m^5^U34 modification improves mitochondrial respiration, as shown by over-expression of the mt-MTO1 enzyme that installs taurine to the modification^59^ in myoblasts of a patient with the A3243G mutation.^59^ It was also shown in A3243G cybrids that co-express the haplotypic T3290C mutation.^60^ These results further demonstrate the close association of this modification with mitochondrial pathology. To measure the level of 1m^5^U34 in the homoplasmic A3243G cybrids, we performed a CMC (*N*-cyclohexyl-*N*’-b-(4-methyl-morpholinium) ethyl-carbodi-imide)-assisted primer extension assay,^16,57,59^ where CMC was used to chemically modify the taurine moiety in total RNA, generating a covalent adduct that was detectable as an RT stop. We observed a strong AMV RT stop at A35 one nucleotide before the modification in the WT, but a much lower level of the stop in the A3243G variant (Figure 2G, left, Figure S2F), indicating a deficiency of the modification in the variant. This deficiency reduced the 1m^5^U34 level to 48% in the variant relative to the WT (Figure 2G, right), as estimated by the fraction of the extension product that terminated at A35 among all extension products (including those that read through the unmodified U34 and terminated at C32 by the presence of ddGTP). Thus, of the modifications that we examined for the A3243G variant, 1m^5^U34 was the only one that was deficient in the variant, consistent with previous reports.^36,58^

### m^1^G9 is the major driver to the altered structure of the A3243G variant

The distinct gel migration of the A3243G variant, and its loss of stability relative to the WT (Figure 2B, C), suggests that it has an altered structure. We tested whether m^1^G9 was the major driver to the altered structure by isolating the effect of the methylation from other natural modifications in mt-Leu(UAA). This was achieved by comparing the structure of the variant relative to the WT tRNA in the m^1^G9-state vs. the native-state. We chose the SHAPE analysis (selective 2’-hydroxyl acylation analyzed by primer extension)^61,62^ for the comparison, which probes RNA local backbone flexibility, where the conformational dynamics of each nucleotide influences the accessibility to a chemical reagent. Each reactive site is then modified by the chemical reagent, which is detected as an RT stop, presenting the conformational dynamics at single-nucleotide resolution. We used NAI (2-methyl-nicotinic acid imidazolide) as the chemical reagent, which maps base-paired and single-stranded regions of RNA with high accuracy.^63^ The results were analyzed for the differential effect of NAI as the ratio of the chemical reactivity at each nucleotide position of the A3243G variant relative to the WT. A ratio of 1 indicated a similar chemical reactivity of the two at a given position, while a ratio different than 1 indicated that the variant had a different reactivity due to base-pairing or a single-stranded feature that was not present in the WT.

We first probed the m^1^G9-state of the A3243G variant vs. the WT to assess the methylation as the single determinant of each tRNA structure (Figure S3A). The results showed variations of the ratio across the tRNA sequence, most notably at positions 12-16, 23-24 and 27-28 in the D stem-loop and at positions 34-40 (Figure 3A, left). In the tRNA canonical structure, these are in the region encompassing m^1^G9, the A14G substitution, and the upper part of the coaxially stacked D-anticodon stem-loops (Figure 3A, right). We next probed the variant vs. the WT in the native state to assess the effect of m^1^G9 in the presence of all natural modifications. Cybrids were grown for the WT and the variant, treated with NAI, and total RNA of each was isolated and probed by RT stops with a primer specific to mt-Leu(UAA). Across the tRNA sequence, the most prominent differences between the variant and the WT were observed at positions 13-18, 24-27, and at the wobble U34 (Figure 3B, left). These regions in the tRNA canonical structure were also mapped to the D-anticodon stem-loops and to the wobble nucleotide U34 (Figure 3B, right). Thus, the mapping data of the m^1^G9-state of the variant relative to the WT extensively overlapped with that of the native-state (Figure 3A, B). Given that m^1^G9 is the single driver that determines the structure of the WT mt-Leu(UAA) (Figure 2A), this extensive overlap indicates that m^1^G9 is also the major driver that determines the structure of the variant. Additionally, the mapping data show how the variant differed from the WT, primarily in the D stem-loop, into the upper part of the anticodon stem-loop, and the wobble U34. The mapping difference at the wobble U34 is noteworthy, given that the variant is deficient of the 1m^5^U34 modification at that position (Figure 2G). It suggests that the structural differences have prevented the variant from being recognized by the enzyme complex mt-MTO1-mt-GTPBP3, which synthesizes 1m^5^U34.^64^

**Figure 3.**
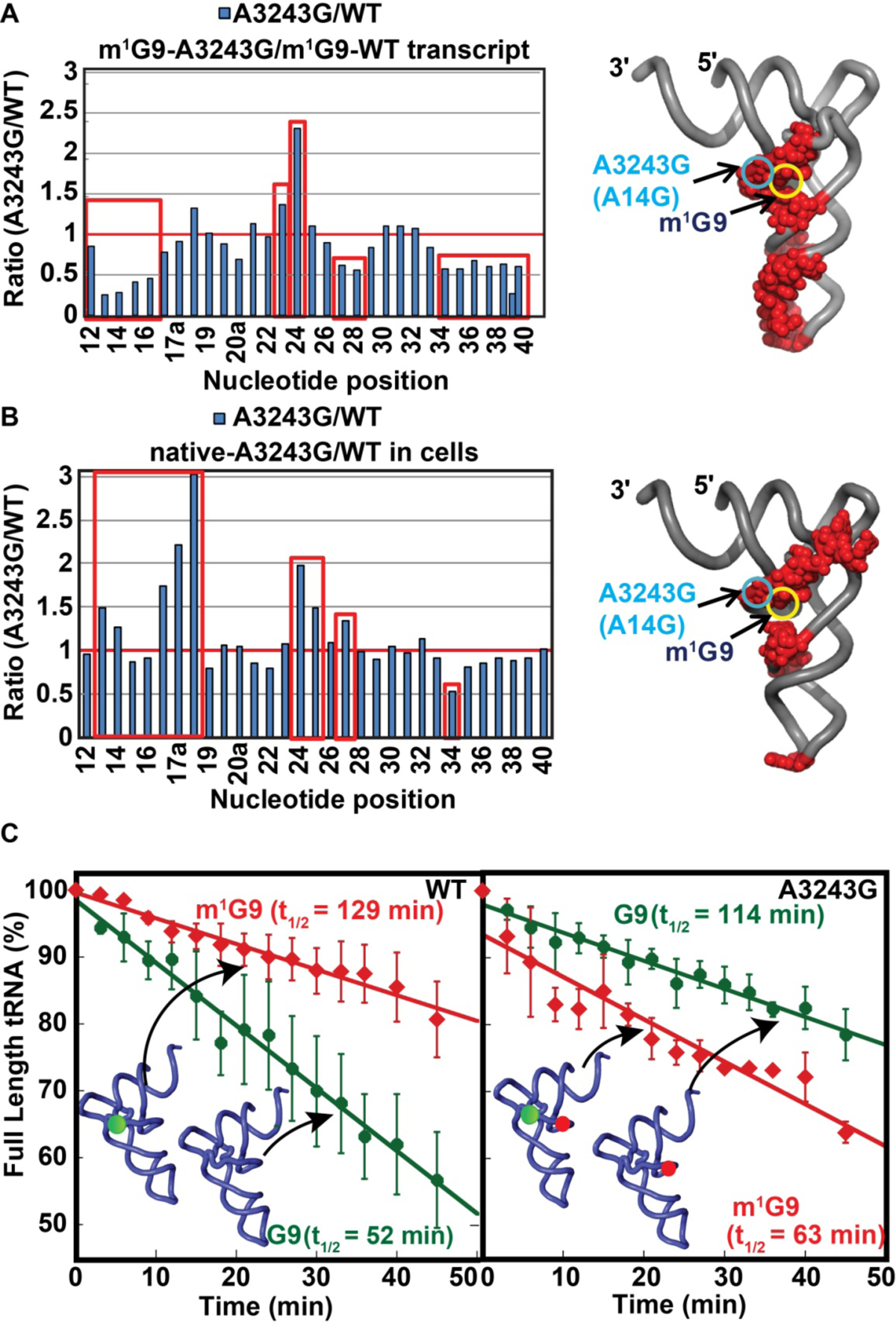
SHAPE and stability analysis of the A3243G variant relative to the WT tRNA (A) SHAPE analysis of the m^1^G9-state of the A3243G variant relative to the m^1^G9-state of the WT tRNA using NAI as the chemical probe. Left: Plots represent the NAI reactivity ratio across nucleotides 12-40, where regions with NAI reactivity above or below the ratio of 1.0 are shown in red boxes. Right: Differential NAI reactivity is mapped in the canonical L-shaped tRNA structure, where the A3243G mutation is shown as a cyan ring, while m^1^G9 is shown as a yellow ring. (B) SHAPE analysis of the native-state of the A3243G variant relative to the native-state of the WT tRNA using NAI as the chemical probe. Left: Plots represent the NAI reactivity ratio across nucleotide positions 12-40, where regions with NAI reactivity above or below the ratio of 1.0 are shown in red boxes. Right: Differential NAI reactivity is mapped in the canonical L-shaped tRNA structure as in (B). (C) Stability of the WT and A3243G variant in the G9-state and the m^1^G9-state. Each transcript was 5’-^32^P-labeled and incubated with a mouse liver mitochondrial lysate over time, where aliquots were analyzed by a 12% PAGE/7M urea sequencing gel. Plots are the average ± SD (n = 3). The half-life t_1/2_ of each tRNA is calculated from curve fitting to a linear regression equation y = *a* + *b*x, where *a* is the dependent variable (intensity at time 0 set to 100), *b* is the slope, y = 50, and x is the half-life t_1/2_.

We then tested whether the m^1^G9-driven variant structure was more sensitive to degradation relative to the WT, based on the notion that RNA in an aberrant structure is usually more prone to degradation.^65^ This would explain the loss of stability of the variant relative to the WT in cybrids (Figure 2C). We prepared G9- or m^1^G9-state of the WT and A3243G variant of mt-Leu(UAA), each 5’-^32^P-labeled, and measured the stability of the label in mitochondrial lysates that were isolated from a mouse liver under conditions that preserved all natural RNases. Due to the use of the label and the focus on m^1^G9 as the single determinant, this stability analysis could not be performed in live cells, where all mt-tRNAs contained the full complement of modifications. Analysis of the label over time showed that, while m^1^G9 as the single modification improved the stability of the WT transcript (t_1/2_ from 52 min of the G9-state to 129 min of the m^1^G9-state), it had an opposite effect on the variant, decreasing its stability (t_1/2_ from 114 min of the G9-state to 63 min of the m^1^G9-state) (Figure 3C, Figure S3C). Thus, m^1^G9 differentially regulates the stability of the WT and the A3243G variant tRNA.

### The folding kinetics of the A3243G and T3271C variants are impeded by m^1^G9

To determine how m^1^G9 guided the A3243G variant to a different and less stable structure than the WT mt-Leu(UAA), we compared their folding kinetics. To isolate the methylation from all other natural modifications in mt-Leu(UAA), we prepared the WT and the A3243G variant in the G9-state and the m^1^G9-state. While the G9-state was made by *in vitro* transcription, the m^1^G9-state was made by an assembly approach, in which the 5’-fragment of each tRNA encoding nucleotides G1 to C17a was chemically synthesized and was joined by T4 Rnl2 with the 3’-fragment encoding nucleotides G18 to A76 generated by *in vitro* transcription (Figure S4A, S4B). We monitored the folding kinetics of each using the enzyme mt-leucyl-tRNA synthetase (mt-LeuRS) to identify the structure that was recognizable for aminoacylation. This previously developed enzyme-based assay^30,66^ used mt-LeuRS as the probe to distinguish between different folding states for different efficiencies of aminoacylation. Each tRNA was incubated without Mg^2+^ and was initiated for the aminoacylation reaction by simultaneous addition of mt-LeuRS and Mg^2+^. In this design, the addition of Mg^2+^ separated the unfolded tRNA species into two populations, one in a meta-stable intermediate-state (due to local intra-molecular base parings) that was competent to fold to the chargeable-state, whereas the other in a disordered state. As shown previously,^30^ the folding of mt-Leu(UAA) from the meta-stable state to the chargeable-state was slow (*k*_obs_ = 0.66 min^-1^), while aminoacylation of the chargeable-state was fast for these enzymes (*k*_obs_ = 10 - 20 s^-1^)^67–71^ (Figure 4A), indicating that the kinetics of aminoacylation was rate-limited by the kinetics of folding from the meta-stable state to the chargeable-state.

**Figure 4.**
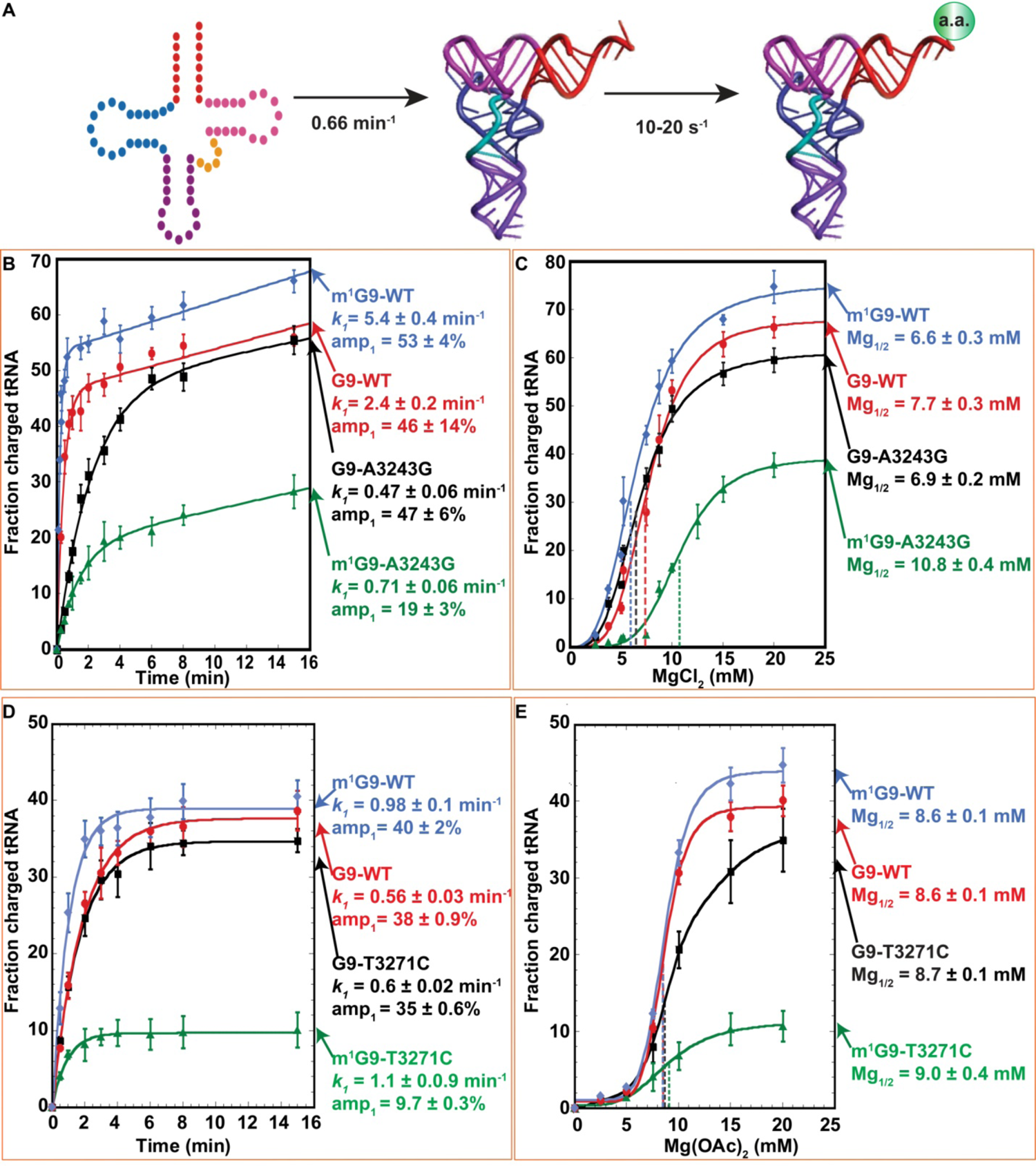
The m^1^G9-state of the A3243G and T3271C variants is impeded for charging (A) An illustration of human mt-Leu(UAA) folding from a meta-stable intermediate-state to the chargeable-state for aminoacylation. The rate constants 0.66 min^-1^ and 10-20 s^-1^ were based on published reports,^30,67–71^ respectively. (B) The fraction of the charged tRNA as a function of time was monitored for the m^1^G9-state of the WT tRNA, the G9-state of the WT tRNA, the G9-state of the A3243G variant, and the m^1^G9-state of the A3243G variant. For each tRNA, the rate constant (*k*_1_) and the amplitude of the first phase (amp_1_) are shown as the average ± SD (n = 3). (C) The fraction of the charged tRNA as a function of MgCl_2_ concentration for the 4 tRNAs described in (B). Each value was the average ± SD (n = 3). (D) The fraction of the charged tRNA as a function of time was monitored for the m^1^G9-state of the WT tRNA, the G9-state of the WT tRNA, the G9-state of the T3271C variant, and the m^1^G9-state of the T3271C variant. For each tRNA, the rate constant (*k*_1_) and the amplitude of the first phase (amp_1_) are shown as the average ± SD (n = 3). (E) The fraction of the charged tRNA at 8 min as a function of Mg(OAc)_2_ concentration for the 4 tRNAs described in (D). Each value was the average ± SD (n = 3).

We observed biphasic kinetics for all 4 tRNAs, where the initial rapid phase represented aminoacylation of the species that were in the meta-stable state and were competent to fold to the chargeable state, while the slower phase represented re-arrangement of the disordered species to the meta-stable state after enzyme addition. For the WT tRNA, the addition of m^1^G9 improved the rate constant *k*_1_ (2.4 ± 0.2 to 5.4 ± 0.4 min^-1^) of aminoacylation of the first phase (Figure 4B), indicating that the methylation facilitated conversion of the meta-stable species to a structure more favorable for charging. The addition of m^1^G9 also increased the amplitude of the first phase (amp_1_) (46 ± 14 to 53 ± 4 %), indicating that it increased the fraction of the meta-stable species to more than 50%. In contrast, the addition of m^1^G9 to the A3243G variant had a negative effect relative to the unmethylated variant. It decreased both *k*_1_ (2.4 ± 0.2 to 0.71 ± 0.06 min^-1^) and amp_1_ (46 ± 14 to 19 ± 3%) (Figure 4B), indicating that the methylation led the meta-stable species to a structure less favorable for charging and that it reduced the fraction of the meta-stable species to less than 20%. As the kinetics continued into the second phase, the most notable deficiency of the m^1^G9-modified A3243G variant was the low level of the aminoacylation amplitude (∼25%), indicating that most of the species were kinetically trapped in a non-chargeable structure. Even the unmethylated variant reached a higher amplitude (∼55%).

The negative effect of m^1^G9 on the A3243G variant was also observed in a titration of Mg^2+^, a diffusible metal ion that can bind discrete folding intermediates.^72^ All 4 tRNAs showed sigmoidal binding kinetics (Figure 4C), indicating that Mg^2+^ binding at one site facilitated binding at another site. In each titration series, the mt-tRNA was incubated with Mg^2+^ for 20 min, after which the aminoacylation reaction was conducted for 3 min, providing information on the fraction of the meta-stable intermediate that was competent to fold into the chargeable state. Notably, the m^1^G9-modified variant exhibited the lowest amplitude (35%) relative to others (60-75%), indicating that it had the lowest fraction that was competent to convert to the chargeable state but the largest fraction that was trapped in a non-competent state. In addition, the m^1^G9-modified variant required the highest Mg^2+^ concentration (10.8 ± 0.4 mM) relative to others to transition from the meta-stable state to the chargeable-state. Even the unmethylated variant required a lower concentration of Mg^2+^ for this transition (6.9 ± 0.2 mM) (Figure 4C).

The results above support a differential effect of m^1^G9 on the folding kinetics of the WT and the A3243G variant. To broaden the scope of this analysis, we tested the T3271C variant, which also contains m^1^G9 in the native state^36^ and represents a distinct structural perturbation relative to the A3243G variant (Figure 1A). As above, the G9- and m^1^G9-state of the WT and the T3271C variant were prepared (Figure S4C). Similar to the A3243G variant, the addition of m^1^G9 to the WT tRNA improved the folding kinetics (*k*_1_ from 0.56 ± 0.03 to 0.98 ± 0.1 min^-1^ and amp_1_ from 38 ± 0.9 to 40 ± 2 %) (Figure 4D), indicating that it led the meta-stable state to a more favorable structure for aminoacylation and that it shifted the fraction of the meta-stable state to a higher level. By contrast, the addition of m^1^G9 to the T3271C variant had a negative effect, substantially reducing the fraction of the meta-stable state (amp_1_ from 35 ± 0.6 to 9.7 ± 0.3%) (Figure 4D). Also, in the Mg^2+^ titration analysis, while the unmodified variant showed similar kinetics as the WT tRNA in reaching a high amplitude (35%), the m^1^G9-modified variant was limited to a much lower amplitude (10%) (Figure 4E), indicating that it was largely trapped in a non-chargeable state. Thus, in both the A3243G and T3271C variants, m^1^G9 impeded the folding kinetics, trapping each in an aberrant and non-chargeable structure. Notably, the kinetic parameters of the A3243G and the T3271C variants are distinct, demonstrating the sensitivity of the kinetic assay to separate the two variants.

### Demethylation of m^1^G9 in cybrids improves the relative stability of the A3243G tRNA

The notion that m^1^G9 has a negative effect on the A3243G variant raised the possibility that demethylation of the variant would improve its structure and activity. We tested this possibility by analysis of the WT and the A3243G variant tRNA in their natural mitochondrial environment to assess the physiological importance of demethylation. Notably, m^1^G9 is synthesized by the ternary complex MRPP1-2-3, where MRPP2 assists the methyl transferase MRPP1 in methylation, while the MRPP1-2 complex assists MRPP3 in 5’-processing of mt-tRNAs^17^ (Figure 5A). Despite the inter-dependence of all three enzymes, each is independently expressed and synthesized in the cytosol and imported into mitochondria. To determine whether demethylation of m^1^G9 from the A3243G variant improved its tRNA structure and activity in mitochondria, we used siRNAs (small interfering RNA) to target *MRPP1* to induce gene-specific KD of the methylation, as there is no small molecular inhibitor of the enzyme. Complete *MRPP1*-KO was not possible, due to the growth-dependent essentiality of the gene.^73,74^ We screened four siRNAs that targeted different regions of the mRNA (Figure S5A, Table S2) and found that 48 h post-transfection had a sufficient KD effect of all four siRNAs on the MRPP1 protein, while 72 h post-transfection had no further effect (Figure S5C, S5D). We monitored 48 h post-transfection and found that siRNA1 and siRNA3 were the most effective in qRT-PCR analysis of the *MRPP1* mRNA, reducing its level to 5% and 12% in WT cybrids, respectively, and to 3% and 15% of the variant, relative to the negative control of each cybrid line mediated by a scrambled siRNA (Figure S5B). These two siRNAs also had the strongest silencing effect on the MRPP1 protein level, reducing it in whole cell lysates of WT cybrids to 22% and 36%, respectively, and to 30% and 34% of A3243G cybrids, in a Western blot analysis (Figure S5C, D). We validated the effect of targeting, showing that the m^1^G9 level in mt-Leu(UAA) was reduced to 85-87% in WT cybrids and to 81-84% in the variant, relative to the negative control and to a non-transfected control in a primer extension assay (Figure 5B).

**Figure 5.**
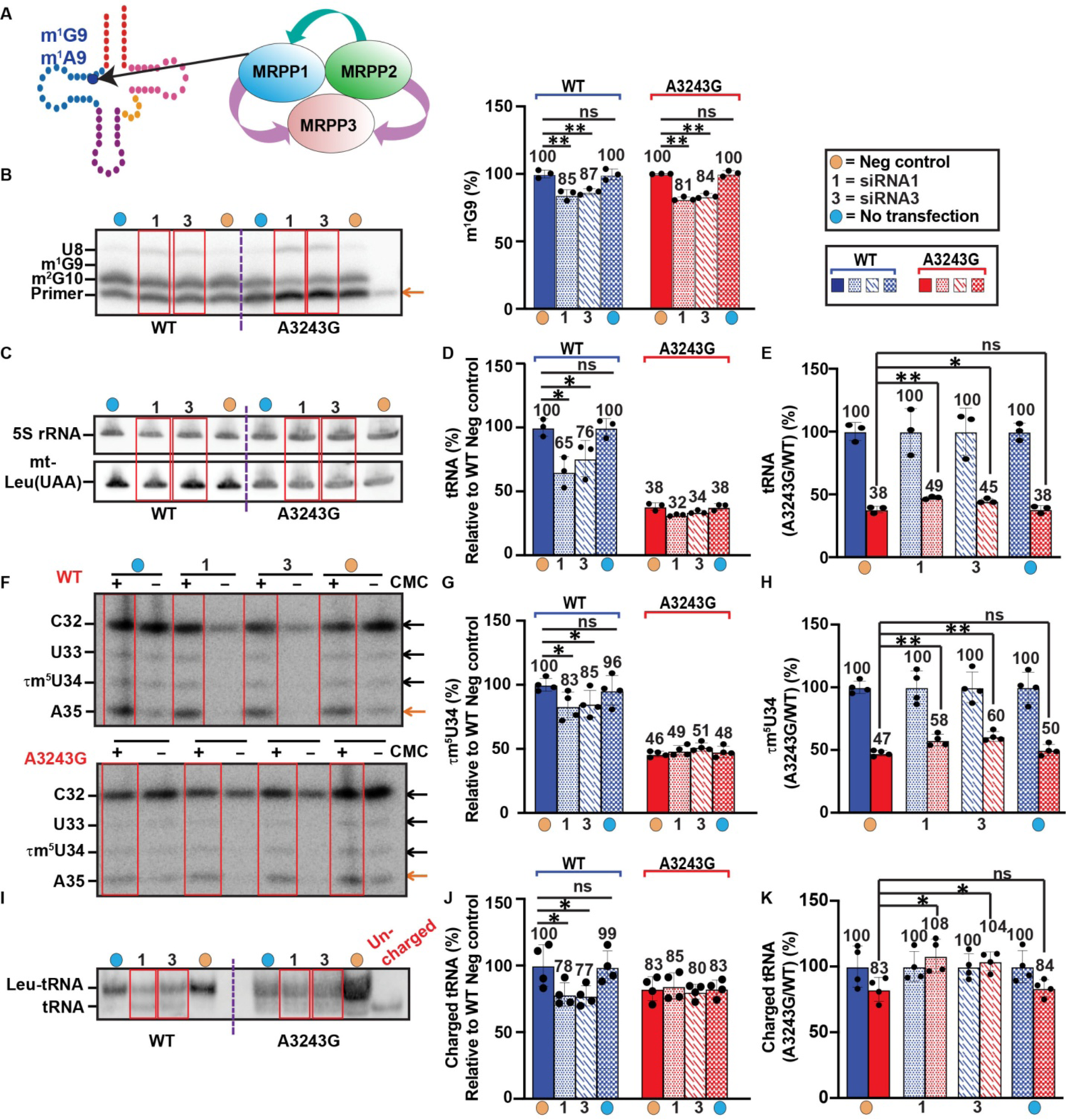
Demethylation of m^1^G9 in mt-Leu(UAA) in cybrids (A) Synthesis of m^1^G9/m^1^A9 in human mt-tRNAs is catalyzed by the MRPP1-2-3 complex, where the methyl transferase MRPP1 is assisted by MRPP2 and the 5’-processing enzyme MRPP3 is assisted by the MRPP1-2 complex. (B) Demethylation of m^1^G9 by siRNA targeting. Left: Demethylation shown in a representative primer extension analysis on total RNA isolated from WT and A3243G cybrids. Yellow dots indicate samples from cybrids transfected with a scrambled siRNA (the Neg control), while blue dots indicate non-transfected samples. All samples were prepared 48 h post-transfection here and below. The m^1^G9 level was calculated by the fraction of the RT stop at position 10 in the sum of positions 10 and 8. Right: Relative m^1^G9 levels to the negative control are shown in the bar graphs, where data are the average ± SD, \*\**p*<0.01 (n = 3). (C) Stability of mt-Leu(UAA) in cybrids analyzed by a representative Northern blot of total RNA of targeted and non-targeted cybrids. (D) Data in (C) shown in a bar graph, where the steady-state level of mt-Leu(UAA) relative to 5S rRNA was quantified for each and normalized to that of WT cybrids in the negative control. Data are the average ± SD, \**p*<0.05 (n = 3). (E) Data in (C) shown in a bar graph, where the steady-state level of mt-Leu(UAA) relative to 5S rRNA was quantified for each and normalized to that of the WT in the same targeting condition. Data are the average ± SD, \*\**p*<0.01, \**p*<0.05 (n = 3). (F) Steady-state levels of 1m^5^U34 analyzed by a representative CMC-assisted primer extension on total RNA of WT and A3243G cybrids. The level of 1m^5^U34 in each sample was calculated as the fraction of A35 in the sum of A35, 1m^5^U34, U33 and C32 and was reported as the level in the CMC (+) lane subtracted from the level of the corresponding CMC (−) lane. (G) Data in (F) shown in a bar graph, where the steady-state level of 1m^5^U34 in mt-Leu(UAA) was quantified for each and normalized to that of WT cybrids in the negative control. Data are the average ± SD, \**p*<0.05 (n = 4). (H) Data in (F) shown in a bar graph, where the steady-state level of 1m^5^U34 in mt-Leu(UAA) was quantified for each and normalized to that of the WT in the same targeting condition. Data are the average ± SD, \*\**p*<0.01 (n = 4). I) Steady-state levels of Leu-tRNA of mt-Leu(UAA) analyzed by a representative acid-urea gel. The level of Leu-tRNA in each sample was calculated as the fraction of the charged and uncharged mt-Leu(UAA). (J) Data in (I) shown in a bar graph, where the steady-state level of Leu-tRNA was quantified for each and normalized to that of WT cybrids in the negative control. Data are the average ± SD, \**p*<0.05 (n = 4). (K) Data in (F) shown in a bar graph, where the steady-state level of Leu-tRNA was quantified for each and normalized to that of the WT in the same targeting condition. Data are the average ± SD, \**p*<0.05 (n = 4). SD is standard deviation and *p* values were obtained from Student’s t test (B, D, E, G, H, J and K). ns is no significance.

Given that the natural m^1^G9 level in the A3243G variant reduced the tRNA stability relative to the WT (Figure 2C), we tested whether a reduction of the methylation by targeting would improve its relative stability. Notably, assessing the effect of demethylation in cells was challenging, because silencing *MRPP1* would reduce the level of m^1^G9/m^1^A9 in all other mt-tRNAs that normally possess the methylation, thus reducing the activity of all of these mt-tRNAs, compromising the mitochondrial activity and ultimately generating an unfavorable environment that negatively affects mt-Leu(UAA). This consideration led us to focus on the relative effect of targeting on the variant vs. the WT mt-Leu(UAA).

Total RNA of the WT and A3243G cybrids were isolated and the level of mt-Leu(UAA) in each was quantified, normalized to the 5S rRNA, and compared to the negative control of the WT as the reference (Figure 5C). This comparison allowed analysis of the effect of targeting on both the WT and the variant at the same time, providing a view into the differential effect. In the negative control, the stability of the variant was 38% relative to the WT (Figure 5D), similar to the value reported above (Figure 2C), whereas upon targeting by siRNA1 and siRNA3, while the stability of the WT substantially decreased from 100% in the control to 65% and 76%, respectively, that of the variant remained unchanged (Figure 5D, Table S3). As a result, comparing the level of each tRNA in the targeted state to the non-targeted control showed that demethylation increased the stability of the variant relative to the WT from 38% to 49% and 45% upon targeting by siRNA1 and siRNA3, respectively (Figure 5E, Table S3).

We assessed whether demethylation of m^1^G9 would affect the 1m^5^U34 modification in the A3243G variant, based on loss of the modification in the non-targeted state (Figure 2G). Total RNA of the WT and A3243G cybrids were isolated and the level of 1m^5^U34 in mt-Leu(UAA) of each cybrid line was probed by the CMC-assisted primer extension assay. While the 1m^5^U34 level of the variant relative to the WT in the negative control was 46% (Figure 5G), similar to that reported above (Figure 2G), this level changed differentially for the WT and the variant upon targeting. For example, upon targeting by siRNA1 and siRNA3, while the level of the WT mt-Leu(UAA) decreased from 100% to 83% and 85%, respectively, that of the variant was unchanged (Figure 5F, G, Table S4). As a result, demethylation increased the 1m^5^U34 level of the variant relative to the WT from 47% to 58% and 60%, respectively (Figure 5H, Table S4). Additionally, we evaluated how demethylation affected the aminoacylation level of the variant. Total RNA of WT and A3243G cybrids were isolated in an acid condition to preserve charged aminoacyl-tRNAs (aa-tRNAs) and were then probed for mt-Leu(UAA) in a Northern blot analysis of an acid-urea gel. Quantification of the charged level of mt-Leu(UAA) in the negative control showed a decrease in the variant to 83% relative to the WT, consistent with previous reports.^36,50^ Upon siRNA1 and siRNA3 targeting, while the charged level in the WT decreased from 100% to 78% and 77%, respectively, that in the variant was unchanged (Figure 5I, 5J, Table S5). As such, demethylation increased aminoacylation of the variant relative to the WT from 83% to 108% and to 104%, respectively (Figure 5K, Table S5). The increases in both 1m^5^U34 and charging in the variant relative to the WT upon targeting suggest an improvement of the variant tRNA structure that is better recognized by the respective enzymes required for the modification and for charging.

### Demethylation of m^1^G9 from the A3243G variant improves mitochondrial respiration

The improvement of the A3243G variant relative to the WT tRNA by demethylation prompted us to test whether it would improve mitochondrial respiration. However, due to the global effect of targeting on all mt-tRNAs that possess the m^1^G9/m^1^A9 methylation, we focused on the early phase of targeting to capture the potential differential effect on cybrids of the WT and A3243G variant. We first assessed the effect of targeting by culturing siRNA-treated cells in a medium containing galactose, where oxidation of the non-fermentable sugar to pyruvate was via glycolysis, yielding no net ATP and forcing cells to generate ATP by mitochondrial oxidative phosphorylation for survival. Thus, when cultured in a medium consisting of galactose as the only carbon source, cells with defective mitochondria would not proliferate. We transfected WT and the A3243G cybrids with siRNA1 or siRNA3 in a glucose medium and then switched them to a galactose medium 48 h post-transfection, which was marked as “Day 0”. Cell counts in the galactose medium over time were then measured relative to Day 0. In WT cybrids, while cells non-targeted or targeted with a scrambled siRNA in the negative control continued to grow in the galactose medium for several days, those targeted by siRNA1 or siRNA3 were immediately arrested in growth (Figure 6A), showing the importance of *MRPP1* for cell survival. In A3243G cybrids, by contrast, while cells non-targeted or targeted in the negative control gradually declined in number over time, indicating a mitochondrial deficiency when cultured in galactose, those targeted by siRNA1 or siRNA3 increased in number from Day 0 to Day 1 and then declined (Figure 6B). Measurement of cell counts relative to WT cybrids in the negative control showed that, while the number of WT declined from 100% to 60% and to 58% upon siRNA1 and siRNA3 targeting, respectively, the cell number of the variant increased from 43% to 90% and 88% upon targeting (Figure 6C), indicating growth. We confirmed that targeting by siRNA1 and siRNA3 was effective even 5 days post-transfection as the MRPP1 protein level remained low relative to 2 days post-transfection (Figure S6A, B). Together, these results showed that demethylation of m^1^G9 by targeting, while destructive to WT cybrids, was beneficial to A3243G cybrids, enabling growth of the latter in the galactose medium.

**Figure 6.**
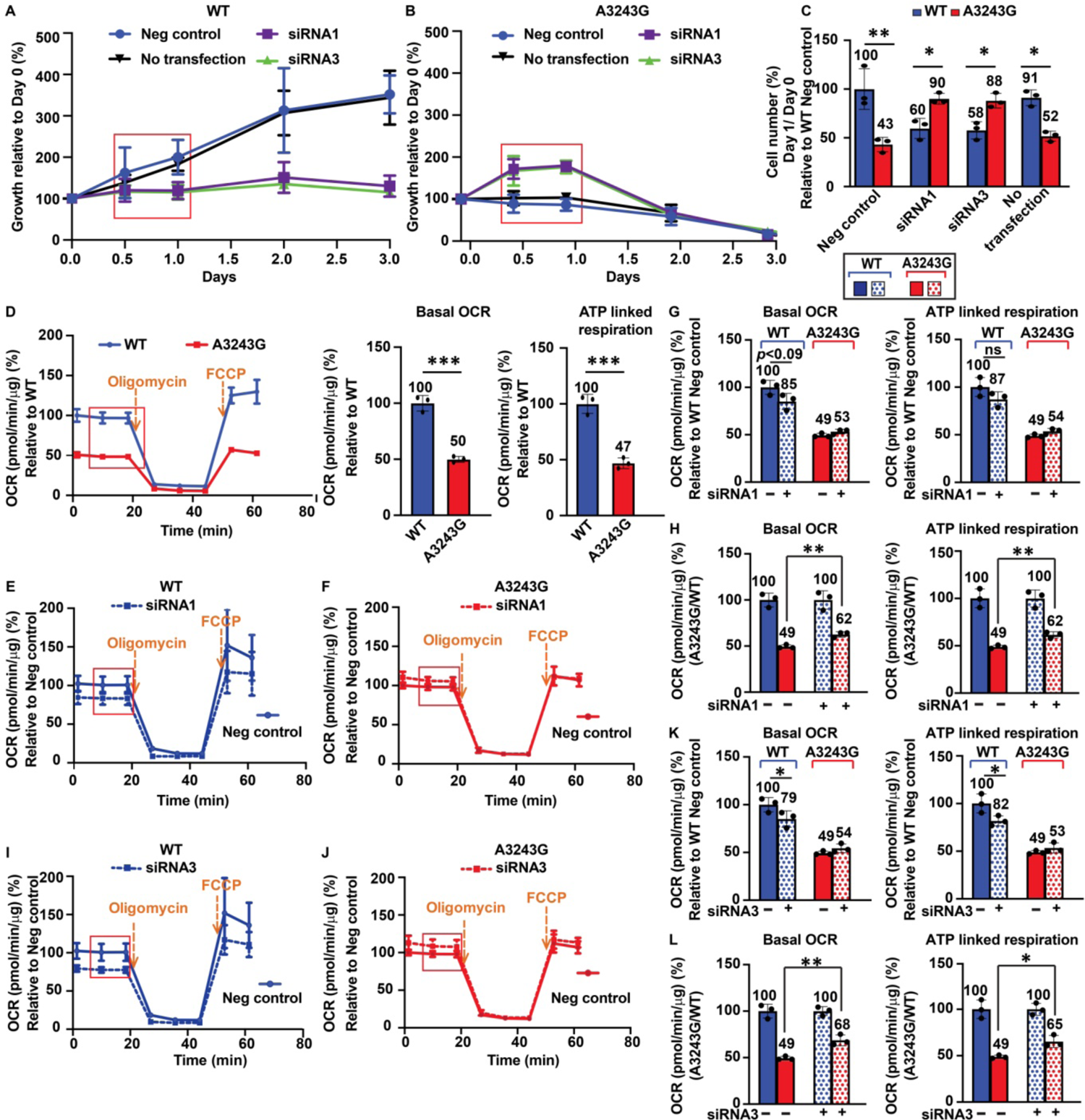
Mitochondrial respiration upon demethylation of m^1^G9 in mt-Leu(UAA) (A) Growth curves in a galactose medium of WT cybrids in different targeting conditions. The region of interest is boxed in red, and data are the average ± SD (n = 3). (B) Growth curves in a galactose medium of A3243G cybrids in different targeting conditions. Data are the average ± SD (n = 3). (C) Bar graphs showing % of cell counts relative to that of WT in the negative control in the region of interest. Each comparison is the average ± SD, \**p*<0.05, \*\**p*<0.01 (n = 3). (D) Measurement of OCR of cybrids with no-transfection. Left: A representative measurement; Middle, Basal OCR of A3243G relative to WT; and Right: ATP-linked respiration of A3243G relative to WT. Data are the average ± SD, \*\*\**p*<0.001 (n = 3). (E) Measurement of OCR of WT upon siRNA1 targeting, showing average ± SD (n = 3). (F) Measurement of OCR of A3243G upon siRNA1 targeting, showing average ± SD (n = 3). (G) Basal OCR and ATP-linked respiration upon siRNA1 targeting of cybrids of A3243G relative to the WT in the negative control, showing average ± SD, *p*<0.09 (n = 3). (H) Basal OCR and ATP-linked respiration upon siRNA1 targeting of cybrids of A3243G relative to the WT in the same targeting condition, showing average ± SD, \*\**p*<0.01 (n = 3). (I) Measurement of OCR of WT upon siRNA3 targeting, showing average ± SD (n = 3). (J) Measurement of OCR of A3243G upon siRNA3 targeting, showing average ± SD (n = 3). (K) Basal OCR and ATP-linked respiration upon siRNA3 targeting of cybrids of A3243G relative to the WT in the negative control, showing average ± SD, \**p*<0.05 (n = 3). (L) Basal OCR and ATP-linked respiration upon siRNA3 targeting of cybrids of A3243G relative to the WT in the same targeting condition, showing average ± SD, \*\**p*<0.01, \**p*<0.05 (n = 3). SD is standard deviation and *p* values were obtained from Student’s t test (C, D, G, H, K and L). ns is no significance.

We next examined the effect of targeting on mitochondrial respiration. We measured the basal oxygen consumption rate (OCR), followed by addition of oligomycin, an inhibitor of ATP synthase, to measure the ATP-linked respiration. This was followed by addition of the uncoupler FCCP (carbonyl cyanide-p-tri-fluoro-methoxy-phenylhydrazone), which disrupts the mitochondrial membrane potential to reveal the maximal electron transport chain activity. Without targeting, A3243G cybrids had 50% of the basal OCR and 47% of the ATP-linked respiration of the WT (Figure 6D), indicating a mitochondrial deficiency. Upon targeting by siRNA1, while WT cybrids lowered both basal OCR and ATP-linked respiration relative to the WT negative control (Figure 6E), the A3243G variant maintained OCR of both (Figure 6F). This contrast was shown in the WT as a reduction from 100% to 85% in the basal OCR and to 87% in the ATP-linked respiration, but no reduction in the variant (Figure 6G). When the respiration of the variant in the targeted state was compared to the WT in the targeted state, we observed an improvement of basal OCR and ATP-linked respiration of the variant from 49% to 62% (Figure 6H). Similarly, upon targeting by siRNA3, while WT cybrids lowered both the basal and ATP-linked respiration relative to the negative control (Figure 6I), the A3243G variant maintained OCR of both (Figure 6J). This contrast was shown in the WT as a reduction from 100% to 79% in the basal OCR and to 82% in the ATP-linked respiration, but no reduction in the variant (Figure 6K). When the respiration of the variant in the targeted state was compared to the WT in the targeted state, we observed an increase in these rates from 49% to 68% and to 65% in the variant (Figure 6L), showing an improvement of mitochondrial respiration.

## Discussion

The inherent structural fragility of mt-tRNAs presents a challenge to understand why these tRNAs are frequently associated with mitochondrial pathology. The discovery that mt-tRNAs are installed with m^1^G9/m^1^A9 during their 5’-processing^17^ suggests that the methylation is the first modification that guides each fragile structure to a stable form.^21^ While this notion emphasizes the need to investigate the role of m^1^G9/m^1^A9 in the folding of mt-tRNAs, it also raises the question of whether the potentially beneficial effect of the methylation on the WT sequence of an mt-tRNA will have the same beneficial effect on its pathogenic variants. However, while the ability of m^1^A9 to guide mt-Lys(UUU) to a native structure is known,^22,23^ specifically only in the WT sequence, how m^1^G9/m^1^A9 affects the folding of other mt-tRNAs has not been studied, particularly in the context of a disease-associated variant. Answering this question will address an unmet need and pave the way to therapeutics for mt-tRNA-associated diseases.

Here we address this unmet need, using MELAS as a model, which represents the most common pathology of all mt-tRNA-associated human diseases.^25^ We focus on the two most frequent mutations of MELAS – A3243G and T3271C in human mt-Leu(UAA), and determine how the m^1^G9 methylation in the tRNA contributes to the folding of the WT and variant tRNAs. We find that, while m^1^G9 indeed promotes stable folding of the WT sequence of mt-Leu(UAA) to a functional structure, it has an opposite effect on the two pathogenic variants, trapping each in an aberrant structure with reduced activity (Figure 7). This is a previously unrecognized notion. Additionally, the parameters and profiles of the folding kinetics differ between the two variants (Figure 4B-E), indicating that each has a distinct aberrant structure. This is consistent with the distinct localizations of the two mutations (Figure 1A), where the A3243G substitution disrupts a loop-stem and loop-turn interaction in the tRNA, while the T3271C substitution disrupts a base-pairing interaction in the anticodon stem. Importantly, while the unmethylated-state of each variant differs from each other in the folding kinetics (e.g., amp_1_ of the G9-state of A3243G at 47 ± 6%, amp_1_ of the G9-state of T3271C at 35 ± 0.6%, a 1.3-fold difference), the m^1^G9-methylated state of each further exacerbates the difference (amp_1_ of the m^1^G9-state of A3243G at 19 ± 3%, and amp_1_ of the m^1^G9-state of T3271C at 9.7 ± 0.3%, a 2.0-fold difference) (Figure 4B, C). This observation indicates that the methylation has an active role in accelerating the decline of each variant, sending each into its aberrant structure. In this parallel analysis of m^1^G9 in two distinct variants of mt-Leu(UAA), while the methylation has a different structural effect on each variant, it has the same trend of taking each variant to a deteriorating structure. This is an important demonstration of the differential effect of m^1^G9 that strengthens conclusions.

**Figure 7.**
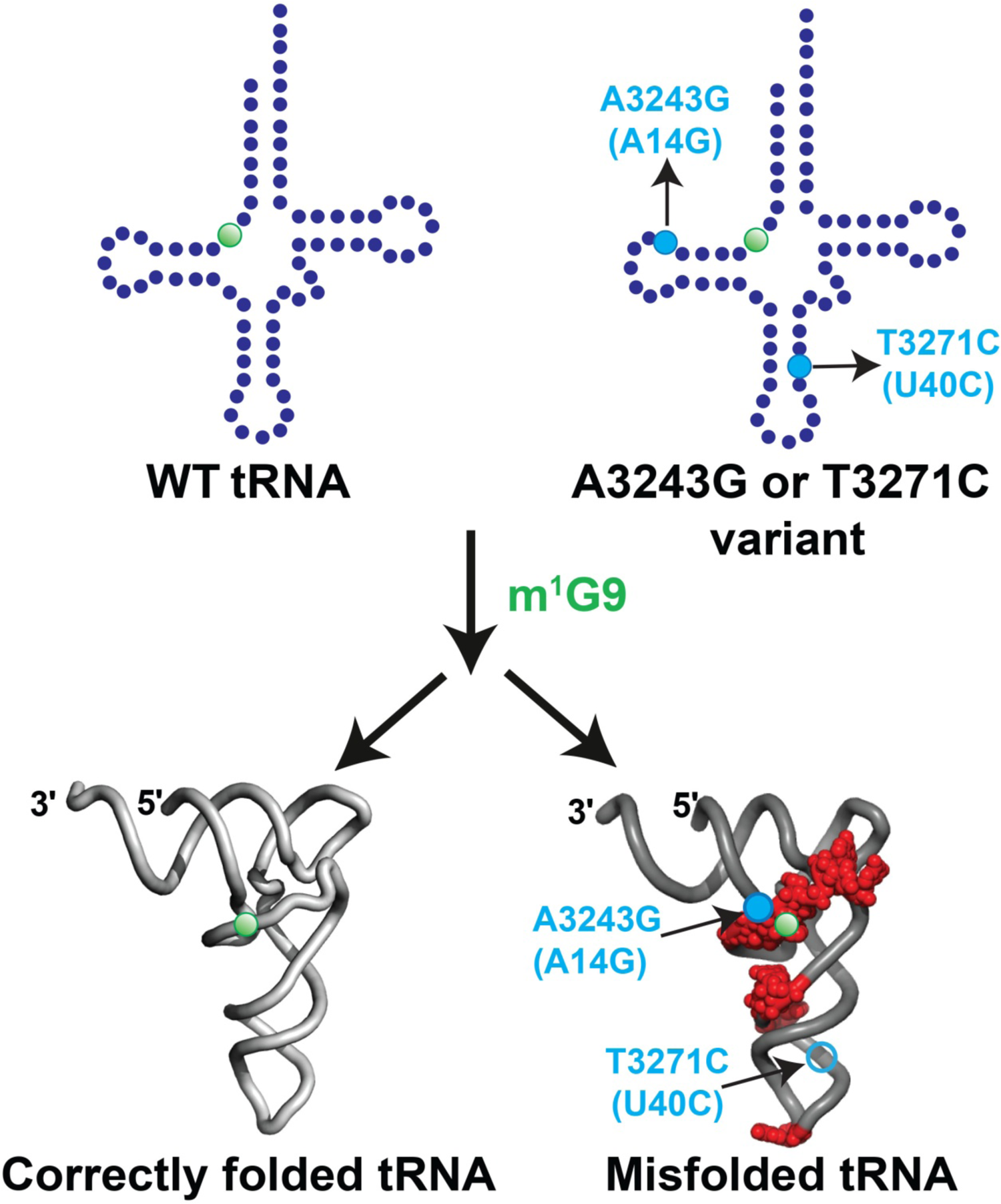
Differential contribution of m^1^G9 to the folding of WT and variants of mt-Leu(UAA) Model for differential contribution of m^1^G9 to the WT and variants of mt-Leu(UAA), represented by the A3243G and T3271C variants. While the addition of m^1^G9 drives the WT tRNA to a correctly folded structure, it drives the A3243G variant to a mis-folded structure that differs from the correct structure in regions marked by red (closed circle). This model is also supported by the folding kinetics of the T3271C variant, although its structural variation has not been mapped (open circle).

The notion that m^1^G9 differentially regulates the functional state of the WT and pathogenic variants of mt-Leu(UAA) is likely generalizable to other mt-tRNA-associated diseases. All 22 human mt-tRNAs have been associated with pathogenic mutations, each of which is mapped to a position that would perturb the tRNA structure (MITOMAP). A recent study of 11 of these pathogenic mutations, localized in various positions throughout sequences of human mt-Ile(GAU), mt-Met(CAU), and mt-Leu(UAA), showed that all had reduced kinetics in the 5’-processing catalyzed by the MRPP1-2-3 complex.^75^ This consistency in the reduction of kinetics indicates that each mutation has resulted in an altered mt-tRNA structure that compromises the 5’-processing activity. Separately, in the known examples of pathogenic variants of mt-tRNAs that have been analyzed for post-transcriptional modifications, all have retained the m^1^G9/m^1^A9 methylation in high stoichiometry (e.g., A3243G, T3271C, and A8344G).^36,51^ Thus, given that the 11 tested pathogenic variants consistently show a destabilizing effect,^75^ and that the characterized pathogenic variants all show a high stoichiometry of the m^1^G9/m^1^A9 methylation,^36,51^ the differential regulation of the methylation in pathology as we report here is an important consideration to understand each disease. Most importantly, our finding that the m^1^G9 methylation exacerbates the destabilizing effects on both the A3243G and T3271C variants (Figure 4B, D) suggests that the methylation-accelerated loss of stability is the root cause of mitochondrial pathology. This accelerated loss of stability would rapidly sensitize each variant to degradation and impair global mitochondrial protein synthesis and respiration. We suggest that this accelerated loss of stability is a factor that precedes the loss of aminoacylation or the loss of the 1m^5^U34 modification of each variant.

The differential effect of m^1^G9 on the WT and the A3243G variant is further supported by analysis of demethylation. In assays for the structure and activity of the A3243G variant relative to the WT mt-Leu(UAA), and in assays for mitochondrial respiration of the variant relative to the WT cybrids, while demethylation is damaging to the WT, it is beneficial to the variant. In all cases, while demethylation reduces the WT activity, it does not change the variant activity, leading to a relative improvement of the variant as compared to the WT upon demethylation (Figures 5, 6). The lack of a robust gain-of-function phenotype of the variant is due to our use of siRNA targeting of *MRPP1* as the means of demethylation, which will remove the methylation in all mt-tRNAs that naturally possess m^1^G9/m^1^A9, thus globally compromising mitochondrial biogenesis. This demonstrates the biological complexity of the demethylation effect.

We suggest new therapeutics for the A3243G-associated MELAS. Instead of using m^1^G9 to strengthen the stability of the variant tRNA, we propose the opposite – removing the methylation to stabilize the variant. The key, however, is to demethylate m^1^G9 only in the A3243G variant, while maintaining the normal level of the methylation in all other mt-tRNAs. Additionally, given the inter-dependence of all three enzymes of the MRPP1-2-3 complex, demethylation of m^1^G9 should be designed not to affect the 5’-processing of all mt-tRNAs, including the A3243G variant, to allow the variant to properly function on the mt-ribosome. This can be achieved by screening for small molecular inhibitors that target the A3243G variant in a complex with MRPP1-2-3 enzymes. As each variant has a unique sequence and structure, as shown in the distinct kinetic profile of the A3243G vs. T3271C variant (Figure 4B-E), this approach is feasible. Indeed, the recent cryo-EM structure of the human MRPP1-2-3 complex, bound with a pre-mt-tRNA substrate,^19^ reveals that the subcomplex MRPP1-2 is sufficient to stably bind to the pre-mt-tRNA substrate and catalyze methylation, which is followed by recruitment of the MRPP3 subunit using a distinct protein-protein interface than the one for methylation. This separation of the 5’-processing activity from the methylation activity is consistent with biochemical analysis^76^ and is attractive for the proposed therapeutic approach. Once small-molecular inhibitors are found from a screening campaign, they can be optimized based on the cryo-EM structure of the pre-mt-tRNA-bound MRPP1-2-3 complex.^19^ Notably, an earlier study showed that over-expression of mt-LeuRS restored mitochondrial respiration of the A3243G variant,^77,78^ while a more recent study showed that over-expression of mt-MTO1 to elevate the taurine level in 1m^5^U34 had the same restorative effect.^59^ However, as the restoration in both cases is mediated by over-expression of an enzyme, it is technically more difficult to translate the result from bench to bedside as compared to small molecular targeting of a site-specific m^1^G9/m^1^A9.

In summary, this work presents a new conceptual framework for understanding how a broadly conserved *N*^1^-methylation of the purine 9 nucleotide in human mt-tRNAs can differentially regulate the fate of a WT mt-tRNA and its pathogenic variants. This new framework changes the course of our conventional thinking of therapeutic development. It also lays the foundation for a new era of research to understand the biological complexity of mt-tRNA-associated pathology.

### Limitations of the study

This work identifies a differential effect of m^1^G9 on the WT mt-Leu(UAA) and its two pathogenic variants of MELAS (A3243G and T3271C), showing that while the methylation is beneficial to the WT tRNA, it is damaging to the variants. While the differential effect was robust in biochemical assays with purified components, it is less robust in cell-based assays. This is due to the use of siRNA targeting in our cell-based assays, which was not specific to mt-Leu(UAA) but would have a general effect on all mt-tRNAs that possess the methylation. Thus, the interpretation of siRNA targeting is limited. Additionally, all cell-based assays were performed using cybrids derived from osteosarcoma cells in this study. These are cells that are not strongly dependent on mitochondrial biogenesis for activity, in contrast to muscle or brain cells. Future experiments are aimed to test the differential effect on patient-derived cardiomyocytes.

## SUPPLEMENTAL INFORMATION

Supplemental Information includes six figures and can be found with this article online.

## ACKNOWLEDGMENTS

We thank Piotr Kopinski of the Wallace lab for sharing the heteroplasmy series of A3243G cybrids, Megumi Hamasaki and Tom Christian of the Hou lab for contributing to the early phase of the project, and all members of the Hou lab for discussion. This work was supported by National Institutes of Health grants R35 GM134931 to Y.M.H., R35 GM149336 to J.P., R01 GM123771 to E.S., CA259635 and AG078814 to D.C.W., and by Japan Society for the Promotion of Science (JSPS 18H05272) and Exploratory Research for Advanced Technology (ERATO, JPMJER 2002) from the Japan Science and Technology Agency (JST) to T.S.

## AUTHOR CONTRIBUTIONS

S.M. designed and performed experiments on most of the tRNA-based assays *in vitro* and *in vivo*, including those for analysis of primer extension, stability, acid-urea gels, Northern blots, and Western blots with and without targeting by siRNAs, while H.G. designed and performed experiments on aminoacylation-based folding assays. S.M. and Y.Y. designed and performed experiments on cell growth assays in galactose. R.Y.H. and M.W. designed and performed experiments on nanopore assays, N.S.L and J.A.P. synthesized m^1^G9-containing RNA oligonucleotides, T.S. isolated the bovine liver mt-Leu(UAA), and E.S. guided measurements of mitochondrial respiration assays. M.W., T.S., E.S., and D.C.W. provided discussion, while Y.M.H. wrote the manuscript. All authors approved the manuscript.

## Star Methods

### Key Resources Table

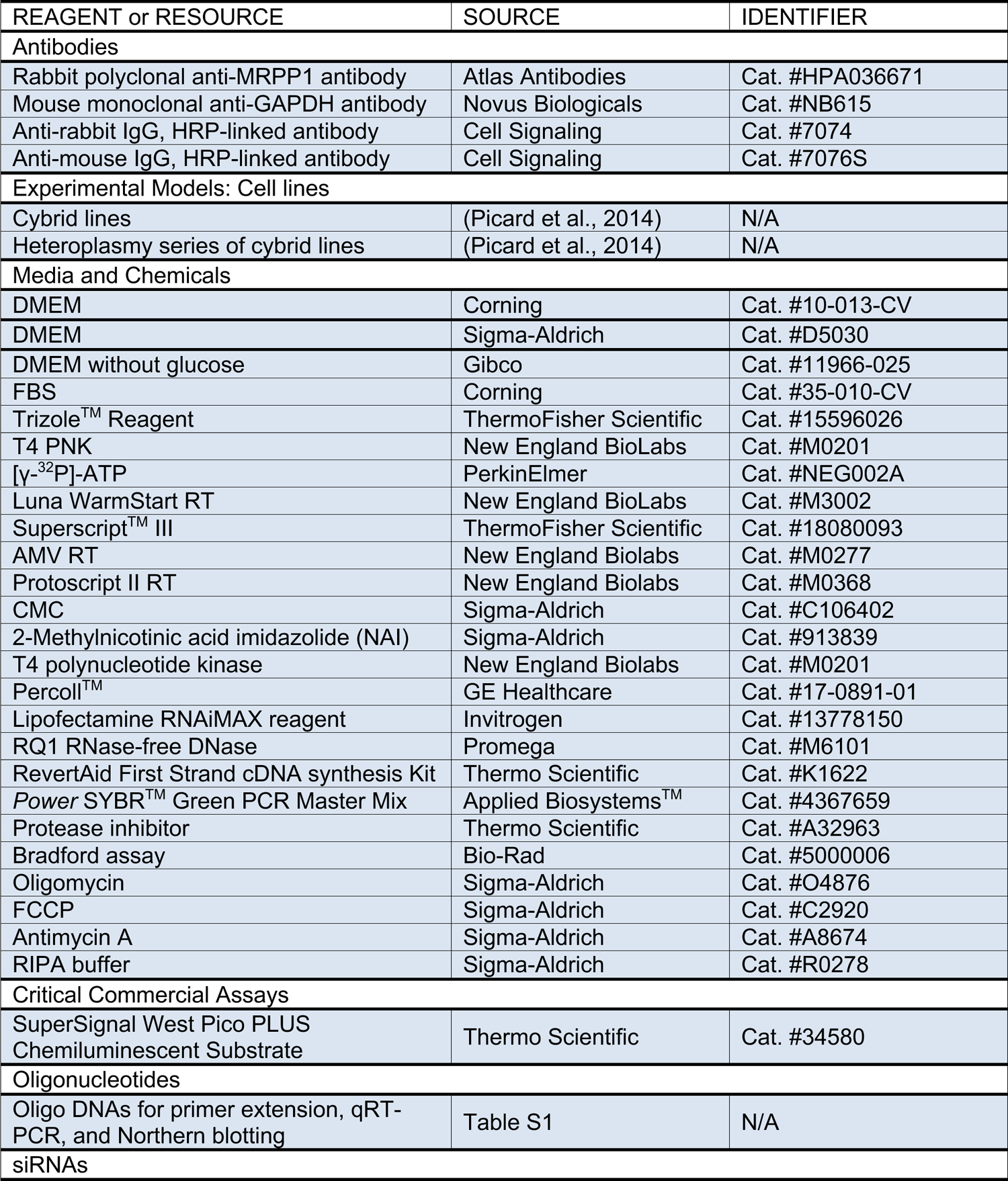
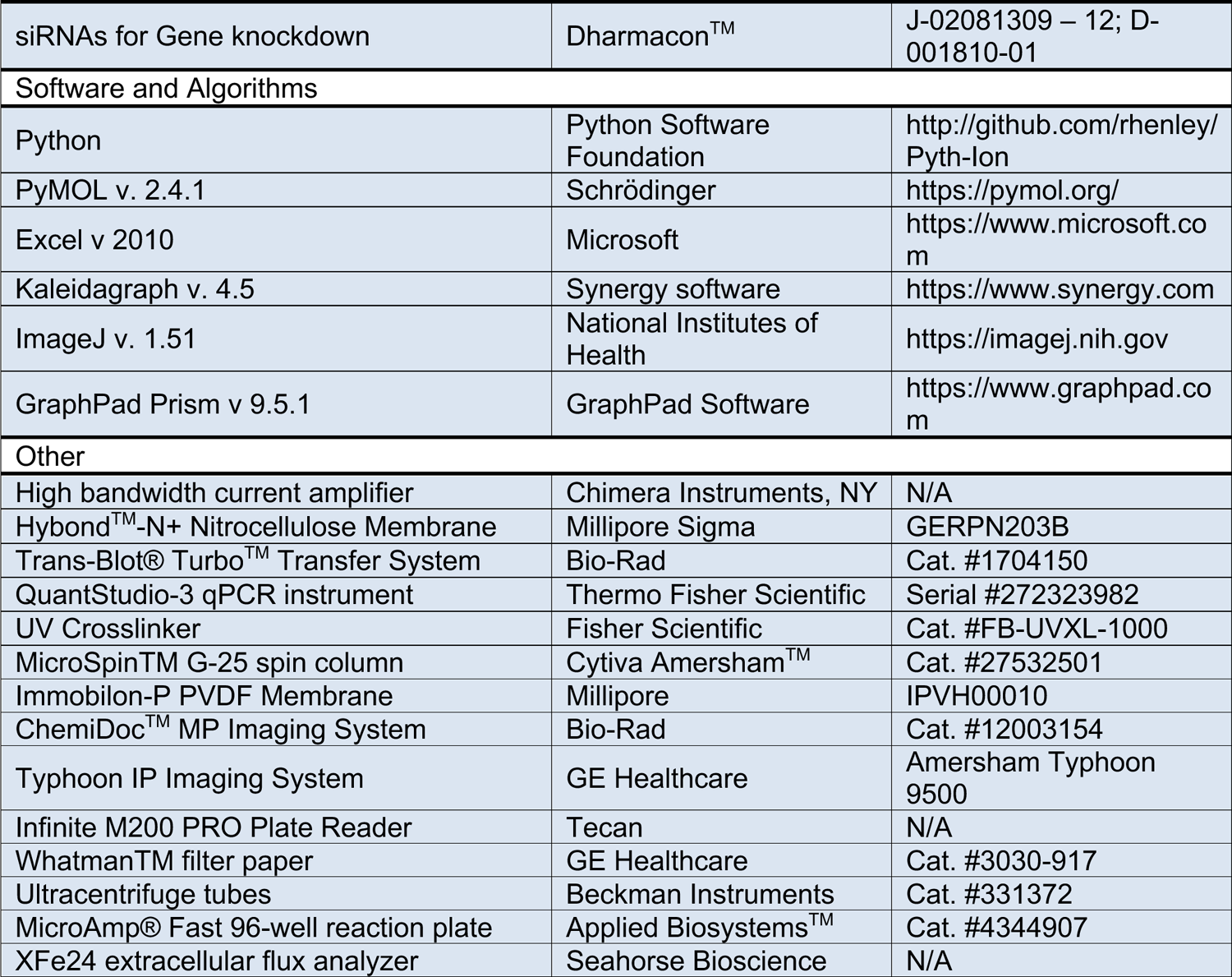

| REAGENT or RESOURCE | SOURCE | IDENTIFIER |
| --- | --- | --- |
| **Antibodies** |  |  |
| Rabbit polyclonal anti-MRPP1 antibody | Atlas Antibodies | Cat. #HPA036671 |
| Mouse monoclonal anti-GAPDH antibody | Novus Biologicals | Cat. #NB615 |
| Anti-rabbit IgG, HRP-linked antibody | Cell Signaling | Cat. #7074 |
| Anti-mouse IgG, HRP-linked antibody | Cell Signaling | Cat. #7076S |
| **Experimental Models: Cell lines** |  |  |
| Cybrid lines | (Picard et al., 2014) | N/A |
| Heteroplasmy series of cybrid lines | (Picard et al., 2014) | N/A |
| **Media and Chemicals** |  |  |
| DMEM | Corning | Cat. #10-013-CV |
| DMEM | Sigma-Aldrich | Cat. #D5030 |
| DMEM without glucose | Gibco | Cat. #11966-025 |
| FBS | Corning | Cat. #35-010-CV |
| Trizole™ Reagent | ThermoFisher Scientific | Cat. #15596026 |
| T4 PNK | New England BioLabs | Cat. #M0201 |
| [γ-<sup>32</sup>P]-ATP | PerkinElmer | Cat. #NEG002A |
| Luna WarmStart RT | New England BioLabs | Cat. #M3002 |
| Superscript™ III | ThermoFisher Scientific | Cat. #18080093 |
| AMV RT | New England Biolabs | Cat. #M0277 |
| Protoscript II RT | New England Biolabs | Cat. #M0368 |
| CMC | Sigma-Aldrich | Cat. #C106402 |
| 2-Methylnicotinic acid imidazolid (NAI) | Sigma-Aldrich | Cat. #913839 |
| T4 polynucleotide kinase | New England Biolabs | Cat. #M0201 |
| Percoll™ | GE Healthcare | Cat. #17-0891-01 |
| Lipofectamine RNAiMAX reagent | Invitrogen | Cat. #13778150 |
| RQ1 RNase-free DNase | Promega | Cat. #M6101 |
| RevertAid First Strand cDNA synthesis Kit | Thermo Scientific | Cat. #K1622 |
| Power SYBR™ Green PCR Master Mix | Applied Biosystems™ | Cat. #4367659 |
| Protease inhibitor | Thermo Scientific | Cat. #A32963 |
| Bradford assay | Bio-Rad | Cat. #5000006 |
| Oligomycin | Sigma-Aldrich | Cat. #O4876 |
| FCCP | Sigma-Aldrich | Cat. #C2920 |
| Antimycin A | Sigma-Aldrich | Cat. #A8674 |
| RIPA buffer | Sigma-Aldrich | Cat. #R0278 |
| **Critical Commercial Assays** |  |  |
| SuperSignal West Pico PLUS Chemiluminescent Substrate | Thermo Scientific | Cat. #34580 |
| **Oligonucleotides** |  |  |
| Oligo DNAs for primer extension, qRT-PCR, and Northern blotting | Table S1 | N/A |
| **siRNAs** |  |  |
| siRNAs for Gene knockdown | Dharmacon™ | J-02081309 – 12; D-001810-01 |
| Software and Algorithms |  |  |
| Python | Python Software Foundation | http://github.com/rhenley/Pyth-Ion |
| PyMOL v. 2.4.1 | Schrödinger | https://pymol.org/ |
| Excel v 2010 | Microsoft | https://www.microsoft.com |
| Kaleidagraph v. 4.5 | Synergy software | https://www.synergy.com |
| ImageJ v. 1.51 | National Institutes of Health | https://imagej.nih.gov |
| GraphPad Prism v 9.5.1 | GraphPad Software | https://www.graphpad.com |
| Other |  |  |
| High bandwidth current amplifier | Chimera Instruments, NY | N/A |
| Hybond™-N+ Nitrocellulose Membrane | Millipore Sigma | GERPN203B |
| Trans-Blot® Turbo™ Transfer System | Bio-Rad | Cat. #1704150 |
| QuantStudio-3 qPCR instrument | Thermo Fisher Scientific | Serial #272323982 |
| UV Crosslinker | Fisher Scientific | Cat. #FB-UVXL-1000 |
| MicroSpin™ G-25 spin column | Cytiva Amersham™ | Cat. #27532501 |
| Immobilon-P PVDF Membrane | Millipore | IPVH00010 |
| ChemiDoc™ MP Imaging System | Bio-Rad | Cat. #12003154 |
| Typhoon IP Imaging System | GE Healthcare | Amersham Typhoon 9500 |
| Infinite M200 PRO Plate Reader | Tecan | N/A |
| Whatman™ filter paper | GE Healthcare | Cat. #3030-917 |
| Ultracentrifuge tubes | Beckman Instruments | Cat. #331372 |
| MicroAmp® Fast 96-well reaction plate | Applied Biosystems™ | Cat. #4344907 |
| XFe24 extracellular flux analyzer | Seahorse Bioscience | N/A |

### RESOURCE AVAILABILITY

#### Lead contact

Further information and requests for resources and reagents should be directed to and will be fulfilled by the lead contact, Ya-Ming Hou.

### Materials availability

Materials generated in this study can be requested from the lead contact.

### Data and Code availability

Information required to reanalyze data reported in this study is available from the lead contact upon request.

## METHOD DETAILS

### Cell culture

WT and cybrid lines with 100% of the A3243G mutation (Picard et al., 2014), were cultured in DMEM (Corning, 10-013-CV) supplemented with 10% FBS (Corning, 35-010-CV), 50 µg/mL uridine and maintained at 37 ⁰C in the presence of 5% CO_2_. Heteroplasmy series of cybrid lines (Picard et al., 2014), with 0, 20, 30, 50, 60, 90 and 100% of the A3243G mutation were cultured under the same condition. Cells were seeded and grown in a fresh culture medium until the confluency was 75%, in 3-4 days, and were split into a fresh medium.

### Nanopore analysis of tRNA folding

All tRNA samples were lyophilized and resuspended in the buffer of 10 mM Tris-HCl, pH 8.0, 1 mM EDTA (TE) to a final concentration of ∼200 ng/µL. For each analysis, an aliquot of each tRNA was diluted to 10 ng/µL in 1X TE with 20 mM MgCl_2_ and incubated at room temperature for ∼30 min. Each was then individually run on a nanopore experimental setup, consisting of a thin insulating SiN membrane that separated two chambers of electrolyte solution (400 mM KCl in 1X TE), where the membrane contained a nanoscale hole of diameter ∼2.7 nm. After incubation, samples were individually added to the *cis* chamber of the setup by replacing the test buffer with sample buffer. A bias voltage of 300 mV was applied across the membrane, driving the negatively charged tRNA molecules in the *cis* chamber through the nanopore in the membrane and towards the positively charged electrode in the *trans* chamber. The current through the nanopore was monitored using a high bandwidth current amplifier (Chimera Instruments, New York, NY), which provided a current sampling rate of 4.16 MHz. This current signal was then filtered at 200 kHz and analyzed using custom python data analysis software (https://github.com/rhenley/Pyth-Ion).

The time-series current data was used to analyze the folding efficiency of each tRNA. The transient current spikes resulting from individual molecules of a tRNA translocating the pore were extracted for the fractional current blockade – the ratio of the signal amplitude to the open pore current value. This fractional current blockade number was fit to a simple geometric model for nanopore conductance to extract a rough size value for the molecule translocating through the pore. Additionally, the dwell time (the duration of the signal; defined as the time for the molecule to translocate the pore) was also determined.

### Northern blot analysis of a heteroplasmy series of A3243G cybrids

A series of cybrid lines with different levels of heteroplasmy of the A3243G mutation (0, 20, 30, 50, 60, 90, and 100%) were cultured in DMEM with 10% FBS, 50 µg/mL uridine and 5% CO_2_ at 37 ⁰C until the cell confluency reached 80% in a 10 cm dish. After removal of the media, the cells were lysed on the plate with addition of 1 mL Trizole^TM^ reagent (ThermoFisher Scientific, 15596026) and processed according to the manufacturer’s protocol to isolate total RNA from each cell line. Each total RNA sample (4 µg) was mixed with 5 µL of a non-denaturing dye (100 mM Tris-HCl, pH 7.0, 50% glycerol, 0.25% xylene cyanol, and 0.25% bromophenol blue), and loaded onto a 12% non-denaturing PAGE in 1X TBE (90 mM Tris-borate, pH 8.0, 2 mM EDTA) buffer in a mini BioRad apparatus, and run for 30 min at 200 V (at 4 ⁰C, cold room). Quantification of the WT and A3243G variant in each cell line was achieved by Northern blot analysis.

Northern blot of each gel, either native or denaturing PAGE, was used to quantify the level of mt-Leu(UAA) in each cell sample. Samples for the native PAGE were prepared as above, while those for the denaturing PAGE were prepared with total RNA (4 μg) from WT and 100% A3243G cells, mixed with 5 µL of a denaturing dye (7 M urea, 0.25% xylene cyanol and 0.25% bromophenol blue), pre-heated at 85 ⁰C for 3 min, and run in a 12% PAGE/7 M urea gel in a mini Bio-Rad apparatus in 1X TBE for 30 min at 200 V (room temperature). Each gel, run in the native or denaturing condition, was electroblotted (25 V and 1 mA for 20 min) under semi-dry conditions onto a positively charged Hybond®-N+ hybridization membrane (Millipore Sigma, GERPN203B) using a Trans-Blot® Turbo^TM^ Transfer System (Biorad, 1704150). The electroblotted membrane was briefly air dried and the RNA was then crosslinked to the membrane using the option of “optimal crosslink” in a UV crosslinker (Fisher Scientific, FB-UVXL-1000). The crosslinked membrane was incubated in a hybridization buffer (0.9 M NaCl, 90 mM Tris-HCl, pH 7.5, 6 mM EDTA, 0.3% SDS, 1% dry milk) at 37 ⁰C for 1 h. A 5’-^32^P-oligonucleotide probe (∼10^6^ cpm), targeting positions 39-56 of human mt-Leu(UAA) (5’-GAACCTCTGACTGTAAAG-3’), was added to the membrane and incubated for 12 h with shaking. Unbound probe was washed off by shaking the membrane twice in 2XSSC buffer (0.3 M NaCl, 30 mM Na-Citrate) for 10-15 min at 37 ⁰C. The membrane was then dried, exposed to an imaging plate overnight, and the tRNA on the membrane was quantified relative to 5S rRNA by a phosphor-imager (Amersham Typhoon model 9500, GE), using ImageJ software (NIH).

### Quantification of modifications by primer extension assays

Cells cultured in a 10 cm dish at 80% confluency were lysed on the plate by replacing the media with 1 mL Trizole^TM^ to isolate total RNA. Primers to monitor and quantify the level of m^1^G9 (5’-GATTACCGGGCTCTG-3’), m^2^G10 (5’-GATTACCGGGCTCT-3’) and m^1^A58 (5’-TGGTGTTAAGAAGAGGA-3’) were synthesized by IDT. Each primer was 5ʹ-labeled with ψ-^32^P-ATP (PerkinElmer) by T4 PNK (polynucleotide kinase, NEB, M0201), purified through a MicroSpin^TM^ G-25 spin column (Cytiva Amersham^TM^, 27532501), and used at 0.33 pmol to anneal with total RNA from WT (3 µg) and with total RNA (6 µg) of MELAS cells in ddH_2_O. The annealing mixture was heated at 90 ⁰C, 3 min, and then slowly cooled to room temperature. To monitor the presence of m^1^G9, Luna WarmStart RT (NEB, M3002) was used for primer extension at 55 ⁰C, 1 h with dCTP (0.5 mM) and ddATP (0.1 mM) in the reaction. To monitor the presence of m^2^G10, Superscript^TM^ III (SSIII) RT (Thermo Fisher Scientific, 18080093) was used for primer extension at 55 ⁰C, 30 min with dGTP (0.5 mM) and ddCTP (0.1 mM) in the reaction. The reaction was terminated by incubation at 70 ⁰C, 15 min. To monitor the presence of m^1^A58, SSIII RT was used for primer extension at 55 ⁰C, 30 min with dATP (0.5 mM) and ddTTP (0.1 mM) in the reaction. The reaction was terminated by incubation at 70 ⁰C, 15 min. An aliquot of each primer extension reaction (5 µL) was mixed with 5 µL denaturing dye, heated at 85 ⁰C, 5 min, and the products resolved on a 12% denaturing PAGE/7 M Urea in 1X TBE in a 40-cm sequencing gel. The gel was transferred to Whatman^TM^ filter paper (GE Healthcare Life Sciences, 3030-917), dried, and phosphorimaged after overnight exposure.

Monitoring of 1m^5^U34 was assisted by pre-treating total RNA with CMC (Millipore Sigma, C106402) as described.^59^ Each RNA sample (15 µg/12 µL of TE) wad diluted with 24 µL of the TEU buffer (50 mM Tris-HCl, pH 8.3, 4 mM EDTA, 7 M Urea), and mixed with 4 µL of CMC (0.1 M final concentration) freshly dissolved in TEU (the CMC reaction) or with 4 µL of TEU (the mocked reaction). The samples were incubated at 30 ⁰C, 16 h, diluted with 160 µL of a stop buffer (50 mM KOAc, 200 mM KCl, pH 7.0), and ethanol precipitated at –80 ⁰C for at least 1 h. The RNA pellet was washed twice with 500 µL of 70% ethanol to remove excess CMC. The pellet was dissolved in 40 µL of an alkaline buffer (50 mM Na_2_CO_3_, 2 mM EDTA, pH 10.4) and incubated at 37 ⁰C, 4 h. Upon dilution to 200 µL with 160 µL of the stop buffer, the RNA was ethanol precipitated as before and dissolved in 10 µL of RNase free water.

The DNA primer (5ʹ-ACCTCTGACTGTAAAG-3ʹ) to detect the CMC-reacted 1m^5^U34 was designed to hybridize to positions 39-54 of the tRNA. It was 5ʹ-labeled with ψ-^32^P-ATP (PerkinElmer), purified through a MicroSpin G-25 column, and used at 0.2 pmol to anneal with the CMC-treated or untreated total RNA (2.5 µg) in 10 µL TE. Annealing was achieved by heating at 80 ⁰C, 2 min, followed by slow cooling to room temperature. AMV RT (NEB, M0277L) was used for primer extension at 42 ⁰C, 1 h, with 1.5 mM each dATP, dTTP and ddGTP. The reaction was terminated by mixing with denaturing loading dye, heat-cooled at 85 ⁰C, 5 min, and the RT products were separated by denaturing 12% PAGE until bromphenol blue reached the bottom of the gel. The level of RT arrest by CMC-modified taurine was quantified by a phosphor-imager as the band intensity that terminated at position 35 relative to the sum of band intensities at positions 32, 33, 34, and 35. To remove background, the ratio obtained from the mock reaction was subtracted from that of the CMC-treated reaction. Each reported value is the average of three independent replicates.

### SHAPE analysis of tRNA structures

For transcripts, 1 µL of 4 µM WT or A3243G variant of mt-Leu(UAA) bearing m^1^G9 was mixed with 16 µL of RNase-free water and heated at 85-90 ⁰C, 3 min. After adding 2 µL of 10X reaction buffer (200 mM MgCl_2_, 400 mM HEPES, pH 7.5), the tRNA was allowed to fold for 15 min at room temperature followed at 4 ⁰C, 15 min. The SHAPE reaction was initiated with addition of 1 µL of 1 M 2-methylnicotinic acid imidazolide (NAI) in DMSO (50 mM final concentration) and continued at 4 ⁰C, 15 min. A control reaction was similarly treated with DMSO alone. Both reactions were stopped by adding DTT to 30 mM final concentration. The tRNA was extracted with phenol-chloroform-isoamyl alcohol, pH 5.0, followed by ethanol precipitation. Each pellet was dissolved in 4.5 µL of RNase free water and half was supplemented with 1 µL of a 5ʹ-^32^P-labeled 21-mer primer (5ʹ-TGGTGTTAAGAAGAGGAATTG-3ʹ; complementary to tRNA positions 56-76) and annealed at 65-70 ⁰C, 2 min, followed by quenching in an ice bath. Primer extension was catalyzed by 50 units of Protoscript II RT (New England Biolabs, M0368) in a 5 µL reaction containing 0.5 mM dNTP, 5 mM DTT, in the 1X Protoscript buffer at room temperature, 5 min, and then 46 ⁰C, 20 min. Reactions were stopped by adding 2.5 µL of denaturing loading dye and heating at 85 ⁰C, 2 min. Aliquots were run through a 12% sequencing gel which was dried and imaged on a phosphor screen after 12-18 h. Bands in the NAI and DMSO lanes were quantified using ImageJ software (NIH). The signal from the DMSO sample was subtracted from the NAI signal to get the value for each NAI reacted band.

For SHAPE analysis of cellular mt-Leu(UAA), WT and A3243G cells grown to 80% confluency were treated with 1 µL of DMSO or 1 µL of 1 M NAI (50 mM final), 15 min. Total RNA was isolated using Trizol^TM^ reagent, 3 µg of which was hybridized to the 5ʹ-^32^P labeled primer as above. All downstream steps were the same as that used for transcript tRNAs.

### Analysis of tRNA stability in mitochondrial lysates

One fresh mouse liver was placed in a beaker to rinse excess blood. Using a razor blade, tissue was thoroughly minced on ice. Minced tissue was transferred to a Potter-Elvehjen homogenization vessel with a Teflon pestle. Homogenization media (30 mL; 250 mM sucrose, 10 mM Tris-HCl, pH 7.4, 0.1 mM EGTA and 0.5% BSA) was added to the vessel and the tissue was gently homogenized using 10 passes of a motorized tissue grinder at a speed of 400 rpm while on ice throughout. The homogenized tissue was transferred to a 30 mL plastic centrifuge tube and was spun in an SS-34 rotor at 2000 rpm at 4 ⁰C, 8 min. The supernatant was transferred to a new tube and was centrifuged at the same speed again. After repeating this step at 8,000 rpm the resulting pellet was resuspended in 1 mL of isolation buffer (250 mM sucrose, 10 mM Tris-HCl, pH 7.4, 0.1 mM EGTA), layered with an additional 14 mL of isolation buffer, and spun at 8,000 rpm 4 ⁰C, 8 min one more time. The pellet was resuspended in 1 mL of mitochondrial resuspension buffer (MRB) (250 mM mannitol, 5 mM HEPES, pH 7.4, and 0.5 mM EGTA) for further processing.

The crude mitochondrial suspension was layered onto 8 mL of a fresh Percoll medium [255 mM mannitol, 25 mM HEPES, pH 7.4, 1 mM EGTA, 30% Percoll (v/v)] followed by 3 mL of MRB in a 13 mL ultracentrifuge tube (Beckman Instruments, Inc, CA 331372) and spun at 95,000 x *g* for 45 min in a TH641 rotor. Mitochondria formed a layer near the bottom of the tube which was carefully pipette out and transferred to a new 13 mL ultracentrifuge tube. After 10-fold dilution with MRB the mitochondria were centrifuged at 6,300 x *g* for 10 min. The mitochondrial pellet was again diluted 1:10 with MRB and centrifuged at 6,300 x *g* for 10 min. The purified mitochondrial pellet was resuspended in 200 µL of the mitochondrial lysis buffer (50 mM Tris-HCl, pH 7.4, 150 mM NaCl, 2 mM EDTA, 2 mM EGTA, 0.2% (v/v) Triton X100, 0.3% NP-40, 0.2 mM PMSF, and a protease inhibitor cocktail) and kept on ice for 30 min to allow mitochondria to lyse. The lysed mitochondria were centrifuged at 12,000 rpm (=13,500 x *g*) for 30 min at 4 ⁰C. The mitochondrial lysate supernatant was transferred to an ice-cold 1.5 mL tube and protein concentration was determined by Bradford.

WT and A3243G transcripts with or without m^1^G9 were used for the *in vitro* stability assay. These four transcripts were 5ʹ-^32^P-labeled and purified in a 40 cm sequencing gel. The excised band of each full-length tRNA was extracted overnight in TE, followed by ethanol precipitation. The purified tRNAs had a specific activity of 10^5^ cpm/µL. Each tRNA was incubated with 2 µL of the mitochondrial lysate (50 µg/µL) at 37 ⁰C in a 50 μL reaction containing 5 µL of 5ʹ-^32^P labeled tRNA (10^4^ cpm/µL) and 20 µL of carrier *E. coli* tRNA (400 ng/µL) in 30 mM HEPES, pH 7.4, 30 mM NaCl, and 5 mM MgCl_2_. Aliquots of 5 µL were removed over time into equal volumes of denaturing gel loading dye. After heating at 85 ⁰C, 3-5 min, each reaction was loaded onto a 12% 40 cm sequencing gel and run at 1,800 V until bromophenol blue reached the bottom. The dried gel was imaged overnight, read on a phosphor-imager, and analyzed using ImageJ software.

### Kinetics of tRNA folding monitored by aminoacylation

Unmodified transcript tRNAs, containing the WT sequence, the A3243G mutation, or the T3271C mutation, were prepared by *in vitro* transcription using in-house T7 RNA polymerase (T7 RNAP). Since the 5’-end of these tRNAs did not contain two consecutive G’s, they were transcribed with an upstream self-cleaving hammerhead ribozyme to generate mature tRNAs with a 5’-hydroxyl group. The m^1^G9-state of each transcript tRNA was prepared by ligation of a chemically synthesized 18-mer 5’-RNA fragment, encoding nucleotides G1 to C17a, with a 60-mer 3’-RNA fragment prepared by *in vitro* transcription, encoding nucleotides G18 to A76. By including GMP in the transcription reaction, most of the RNAs contained a 5’-phosphate. Ligation reactions were carried out in the absence of a splint by incubating equimolar amounts of the 18-mer and the 60-mer RNAs with T4 Rnl2 and ATP at 37 ⁰C for 1 h. All transcripts and ligation products were purified by denaturing 12% PAGE.

Aminoacylation by mt-LeuRS of each transcript tRNA, in the G9-state or the m^1^G9-state, was performed under single-turnover conditions as described.^66^ Each tRNA was 3’-labeled with ^32^P by exchange with α-^32^P ATP in a reaction catalyzed by *Bst1* CCA adding enzyme at 60 ⁰C, 30 min. The labeled tRNA was extracted with phenol-chloroform-isoamyl alcohol, spun through a gel filtration cartridge, and ethanol precipitated. Specific activity of the labelled tRNA was 20,000-100,000 cpm/pmole. Prior to aminoacylation, each labelled tRNA (1 pmol) was incubated 20 min at room temperature in 5 µL of 1X reaction buffer (80 mM HEPES, pH 7.0, 5 mM DTT, and 8 mM leucine). Aminoacylation was initiated by adding an equal volume of 2X assay mix (10 µM LeuRS, 20 mM Mg(OAc)_2_, and 25 mM ATP/Mg in 1X reaction buffer) at room temperature. Aliquots of 1 µL were removed over time into 3 µL of quench solution (50 mM NaCl, 200 mM NaOAc, pH 5.0) and stored at –20 ⁰C until analyzed. Folding as a function of Mg(OAc)_2_ concentration was investigated by incubating each tRNA for 20 min at room temperature with the indicated concentrations of Mg(OAc)_2_ in X1 reaction buffer. Aminoacylation was carried out for 3 min at room temperature as described without changing the Mg(OAc)_2_ concentration. Aminoacylation was determined by treating each quenched aliquot with 6 units of S1 nuclease at 37 ⁰C, 20 min, followed by thin layer chromatography on PEI-cellulose 0.1M NH_4_Cl/5% HOAc. The ratio of labeled AMP to leucyl-AMP was determined by phosphor-imaging.

### Targeting *MRPP1* with siRNAs

Cultured WT and A3243G (100% mutation) lines (60 x 10^3^ cells) were seeded on each well of a 12 well-plate one day before transfection to reach a confluency of 40-50% for transfection.

The cell number was adjusted based on the plate size to 40-50% at the time of siRNA transfection. The cells were transfected with commercially available, predesigned, chemically synthesized siRNAs targeted to the MRPP1 gene (Dharmacon^TM^, J-020813 09-12) along with a scrambled siRNA (Dharmacon^TM^, D-001810-01) as a negative control at 30 nM final concentration using Lipofectamine RNAiMAX reagent (Invitrogen, 13778150), following the manufacturer’s protocol. After 24 h of transfection, culture media were replaced with fresh DMEM/10% FBS. Cells at 48 h post-transfection were harvested for analysis of the mRNA and protein levels of *MRPP1* by qRT-PCR and Western blot, respectively.

The effect of siRNA targeting on the *MRPP1* mRNA was evaluated by qRT-PCR analysis. Total RNA was extracted from transfected cells using TRIzol^TM^ according to the manufacturer’s protocol. Genomic DNA was removed from the total RNA by treatment with RQ1 RNase-free DNase (Promega, M6101). cDNA synthesis was carried out with the RevertAid First Strand cDNA synthesis Kit (Thermo Scientific, K1622). qPCR was performed on a QuantStudio-3 qPCR instrument (Thermo Fisher Scientific). For targeting *MRPP1*, 18 µL of qPCR master mix (10 µL of *Power* SYBR^TM^ Green PCR Master Mix (Applied Biosystems^TM^, 4367659), 7.6 µL of nuclease-free PCR grade water, and 0.4 µL of gene-specific forward and reverse primers (0.2 pmol each)) were combined with 2 µL of the cDNA template into each well of a 96-well reaction plate (MicroAmp® Fast 96-well reaction plate, 0.1 mL, Applied Biosystems^TM^, 4344907). qPCR amplification was performed using the following cycling conditions: preincubation 95 ⁰C, 10 min, followed by 45 cycles of 3 steps of amplification with 95 ⁰C (10 s), 48 ⁰C (10 s), and 72 ⁰C (1 min). The transcript level for each gene was determined using a standard curve generated from the control template by plotting the Cq values (y-axis) against the log initial concentration (x-axis). GAPDH was used for normalization between samples. The primer sets used for amplification of *MRPP1 and GAPDH* were: *MRPP1* forward primer: 5ʹ-GAGTACACCCCCTTCTGAAG-3ʹ; *MRPP1* reverse primer: 5ʹ-GGGTTTTGAGCTCTTCTTCAG-3ʹ; *GAPDH* forward primer: 5ʹ-ATGGGCAAGGTGAAGGTG-3ʹ; *GAPDH* reverse primer: 5ʹ-GTACACCATGTAGTTCAGGTC-3ʹ.

The effect of siRNA targeting on protein expression of MRPP1 was analyzed by Western blot analysis. Transfected cells were washed with ice cold PBS three times and lysed on the plate by adding 1 mL of NP-40 lysis buffer (50 mM Tris-HCl, pH 8.0, 150 mM NaCl, 1% NP-40; supplemented with Protease inhibitor (Thermo Scientific^TM^, A32963)) on an ice bath, 30 min. Supernatant was collected from lysed cells by spinning at 16,000 x *g*, 20 min, and samples were quantified by Bradford assay (Bio Rad, 5000006) for even gel loading. For MRPP1 and GAPDH detection, 20 µg of protein was run on a 12% SDS-PAGE, and then transferred to a PVDF membrane (Millipore, IPVH00010) by semi-dry blotting using the Trans-Blot® Turbo^TM^ Transfer System (BioRad, 1704150). After 1 h blocking with 5% skim milk in TBST (1X TBS, 0.1% Tween-20), the membrane was incubated overnight with anti-MRPP1 antibody (Atlas Antibodies, HPA036671) at 1:1000 dilution or anti-GAPDH antibody (Novus Biologicals, NB615) at 1:2500 dilution at 4 ⁰C with gentle agitation. After three 10 min washes with 1X TBST, the membrane was incubated with anti-rabbit secondary antibody (Cell Signaling, 7074) at 1:2000 dilution for the MRPP1 membrane and anti-mouse secondary antibody (Cell Signaling, 7076S) at 1:5000 dilution for the GAPDH membrane for 1 h at room temperature in 1X TBST, followed by three washes of 10 min each. The membrane was incubated with SuperSignal^TM^ West Pico PLUS chemiluminescent substrate (Thermo Scientific, 34580), 1-2 min, and the signal was detected using the ChemiDoc^TM^ MP Imaging System (Bio-Rad, 12003154). Band intensity was quantified using ImageJ software.

### Growth curve in galactose media

A 48 well plate was seeded with 0.5 x 10^4^ cells per well in 200 µL of glucose media. The cells were transfected with *MRPP1* siRNA1 and siRNA3 along with a scrambled siRNA as a negative control (final siRNA concentration 30 nM) using Lipofectamine RNAiMAX reagent (Invitrogen, 13778-075). Galactose media were prepared by using DMEM without glucose (Gibco, 11966-025) but supplemented with 10 mM galactose, 2 mM glutamine, 5 mM HEPES, 1 mM sodium pyruvate and 10% FBS. Cells were washed with PBS after 48 h of transfection and media were changed to galactose media. The cells in each well were counted after trypsinization every 24 h for 3-5 days unless mentioned otherwise.

### Acid-Urea PAGE analysis of tRNA aminoacylation levels in cybrids

Total RNA was extracted from cells in an acidic condition (pH 4.5) to maintain aa-tRNA levels.^79^ Cells harvested from a 10 cm culture dish (1×10^6^ cells) were mixed at a 1:1 volume ratio with 10% TCA and incubated on ice, 10 min. Cells were spun down at 8,000 rpm (= 7,600 x *g*) at 4 ⁰C, 5 min, resuspended in an extraction buffer (0.3 M NaOAc, pH 4.5, 10 mM EDTA), and mixed with an equal volume of ice-cold phenol-chloroform-isoamyl alcohol (83:17:3.4), pH 5.0. The mixture was vortexed for 5 cycles of 1 min vortex and 1 min rest on ice and spun at 12,000 rpm (= 13,500 x *g*) at 4 ⁰C, 10 min. An equal volume of cold isopropanol was added to the aqueous phase which was incubated at −20 ⁰C, 30 min for precipitation. The nucleic acid pellet was collected after centrifugation at 12,000 rpm (= 13.500 x *g*) at 4 ⁰C, 20 min, washed with 70% ethanol in 30% 10 mM NaOAc, pH 4.5, dried, and dissolved in gel-loading buffer (10 mM NaOAc, pH 4.5,1 mM EDTA). Nucleic acid from WT cells (2.5 µg) and from MELAS cells (5 µg) were mixed with 2 volumes of an acid-urea gel loading buffer (0.1 M NaOAc, pH 5.0, 9 M urea, 0.05% bromophenol blue (BPB), and 0.05% xylene cyanol (XC)). An intermediate sized acid-urea gel (14 x 17 x 0.5 mm) was used to resolve charged from uncharged mt-Leu(UAA). Each acid-urea gel was made with 6.5% PAGE/7 M urea in 0.1 M NaOAc, pH 5.0, and run at 250 V for 3 h 45 min at 4 ⁰C. Following electrophoresis, the region between BPB and XC was excised and washed in 1X TBE. Nucleic acid was transferred from the gel to a wetted nitrocellulose membrane in 1X TBE and quantified by Northern blot analysis using the probe described above. The charged fraction was calculated as the area encompassing the charged band divided by the sum of charged and uncharged bands.

### Measurement of oxygen consumption rate and ATP-linked respiration

The oxygen consumption rate in cybrid cell lines was detected using an XFe24 extracellular flux analyzer (Seahorse Bioscience) as described^59^ with some modifications. Cells that were transfected with MRPP1 siRNAs along with the scrambled siRNA as a negative control were trypsinized and counted 24 h post-transfection. Cells (10^4^ counts) were seeded in 4 wells of a XFe24 cell culture miniplate precoated with collagen (Seahorse Biosciences) and cultured for an additional 24 h to further knockdown *MRPP1*. After 24 h, the culture medium was replaced with freshly resuspended DMEM (Sigma-Aldrich, D5030), pH 7.4, containing 25 mM glucose and 4 mM L-glutamine. The rates of oxygen consumption were measured under basal conditions and over the course of programmed injections of oligomycin (Sigma-Aldrich, O4876) (final concentration 5 µg/µL), carbonyl cyanide-p-tri-fluoro-methoxy-phenylhydrazone (FCCP) (Sigma-Aldrich, C2920) (final concentration 500 nM), and antimycin A (Sigma-Aldrich, A8674) (final concentration 10 µM), which is an antibiotic that specifically inhibits metabolic reduction of cytochrome C. Basal respiration was determined as the OCR before oligomycin, subtracted from the minimum OCR after the antibiotic treatment. The ATP-linked respiration rate was determined as the OCR before oligomycin subtracted from the minimum OCR after oligomycin. After completion of measurements, the cells were lysed with the RIPA buffer (Sigma-Aldrich, R0278) and protein concentration was measured by Bradford assay to normalize the cell number variation in each well.

## QUANTIFICATION AND STATISTICAL ANALYSIS

The statistical significance of biochemical and cell-based assays was tested by Student’s t-test.

## Supplemental Figures

**Figure S1.**
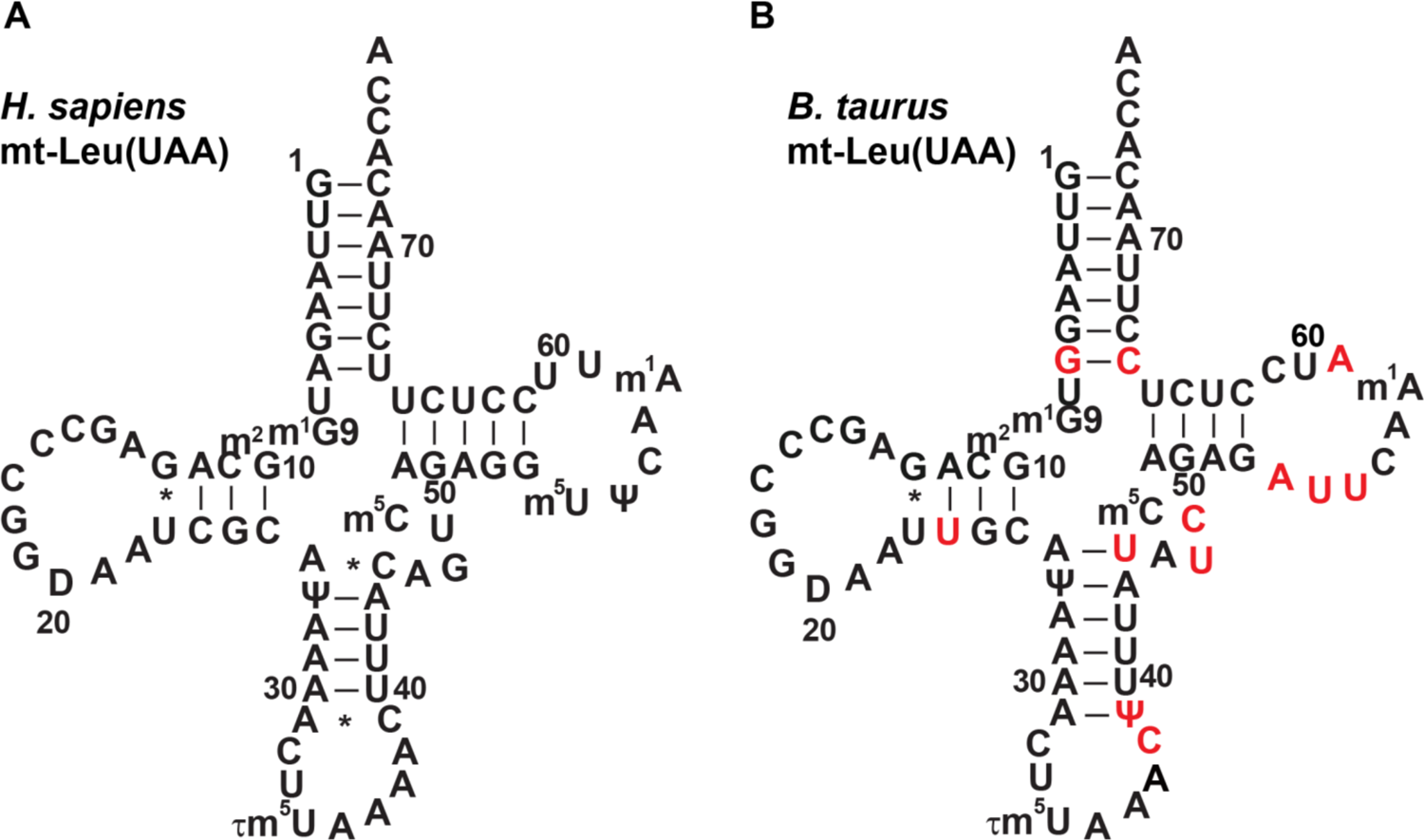
(Related to Figure 1) Comparison of mt-Leu(UAA) of human *Homo sapiens* and bovine *B. taurus*. (A) Sequence and cloverleaf structure of *H. sapiens* mt-Leu(UAA) in the native-state.^16^ (B) Sequence and cloverleaf structure of *B. taurus* mt-Leu(UAA) in the native-state,^41^ where differences from the human counterpart are highlighted in red.

**Figure S2.**
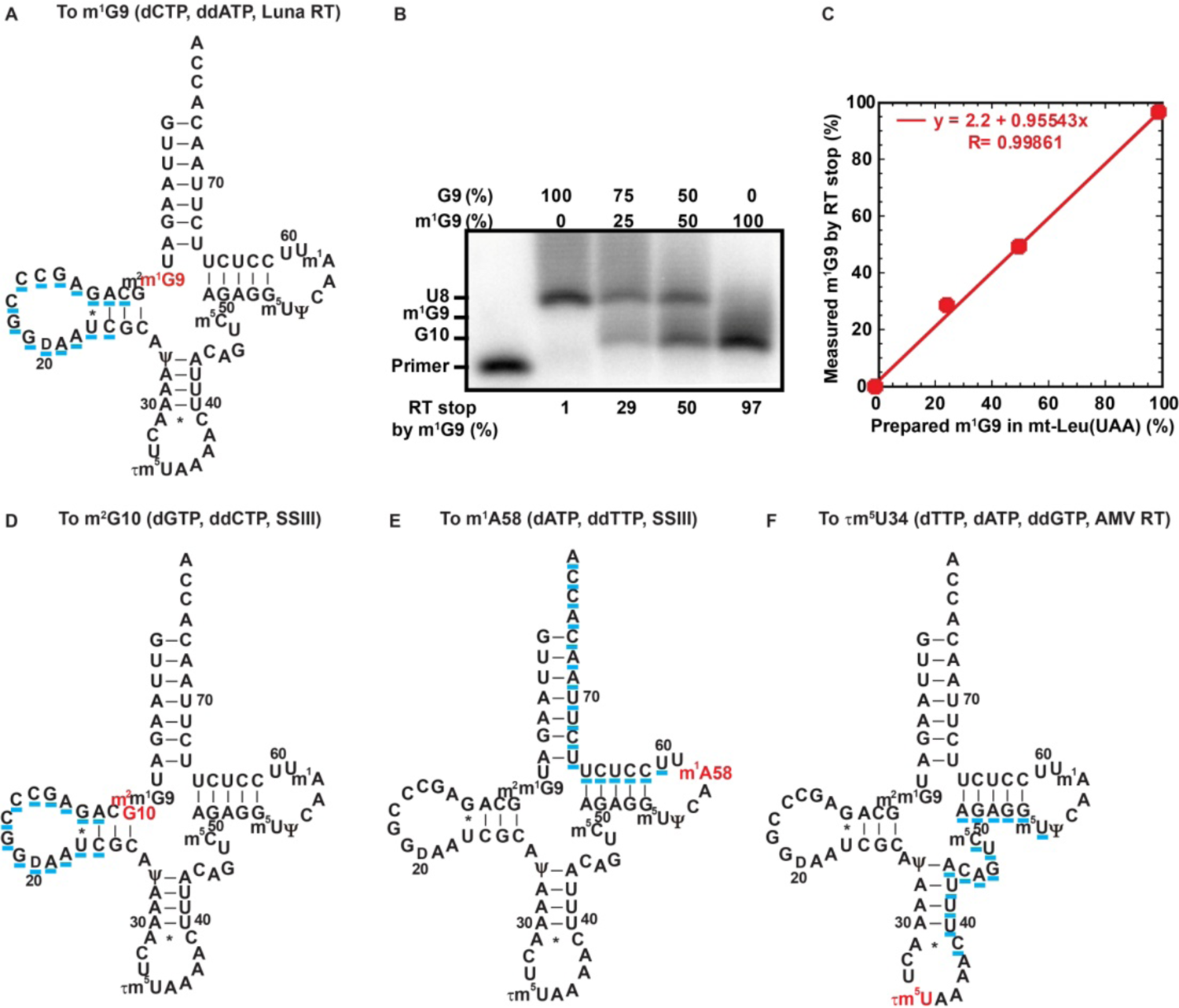
(Related to Figure 2) Design of primer sequences to quantify modifications in human mt-Leu(UAA) by primer extension. (A) The targeted sequence (underlined in blue) in human mt-Leu(UAA) by primer extension to quantify the level of m^1^G9 in a reaction with dCTP, ddATP, and Luna RT. (B) Denaturing gel analysis of the Luna RT stop on the transcript of mt-Leu(UAA) harboring m^1^G9 at 0, 25, 50, and 100%, showing the % of the RT stop relative to the readthrough to U8. (C) A linear correlation between the measured RT stop in (B) and the actual fraction of m^1^G9 in the transcript of mt-Leu(UAA) as prepared. (D) The targeted sequence (underlined in blue) in human mt-Leu(UAA) by primer extension to quantify the level of m^2^G10 in a reaction with dGTP, ddCTP, and SSIII. (E) The targeted sequence (underlined in blue) in human mt-Leu(UAA) by primer extension to quantify the level of m^1^A58 in a reaction with dATP, ddTTP, and SSIII. (F) The targeted sequence (underlined in blue) in human mt-Leu(UAA) by primer extension to quantify the level of 1m^5^U34 in a reaction with dTTP, dATP, ddGTP, and AMV RT.

**Figure S3.**
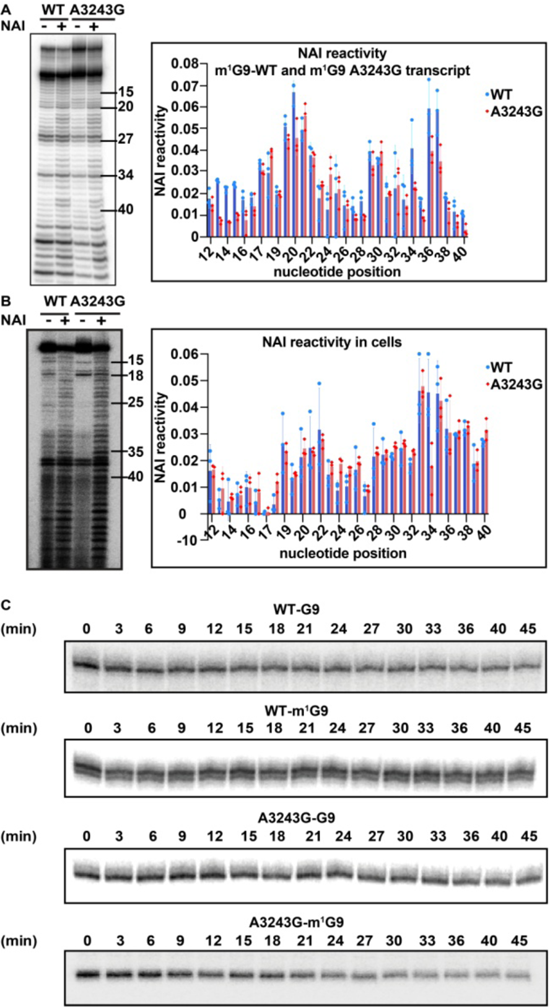
(Related to Figure 3) SHAPE analysis of mt-Leu(UAA) (A) Gel analysis of a representative SHAPE reaction on the m^1^G9-state of WT and A3243G of mt-Leu(UAA). NAI reactivity in WT and A3243G across nucleotides 12-40 is plotted on the right. (B) Gel analysis of a representative SHAPE reaction on the native-state of WT and A3243G of mt-Leu(UAA). NAI reactivity in WT and A3243G across nucleotides 12-40 is plotted on the right. (C) Stability of 5’-^32^P-labeled WT and A3243G variant of mt-Leu(UAA), with and without m^1^G9, over time incubated in a mitochondrial lysate.

**Figure S4.**
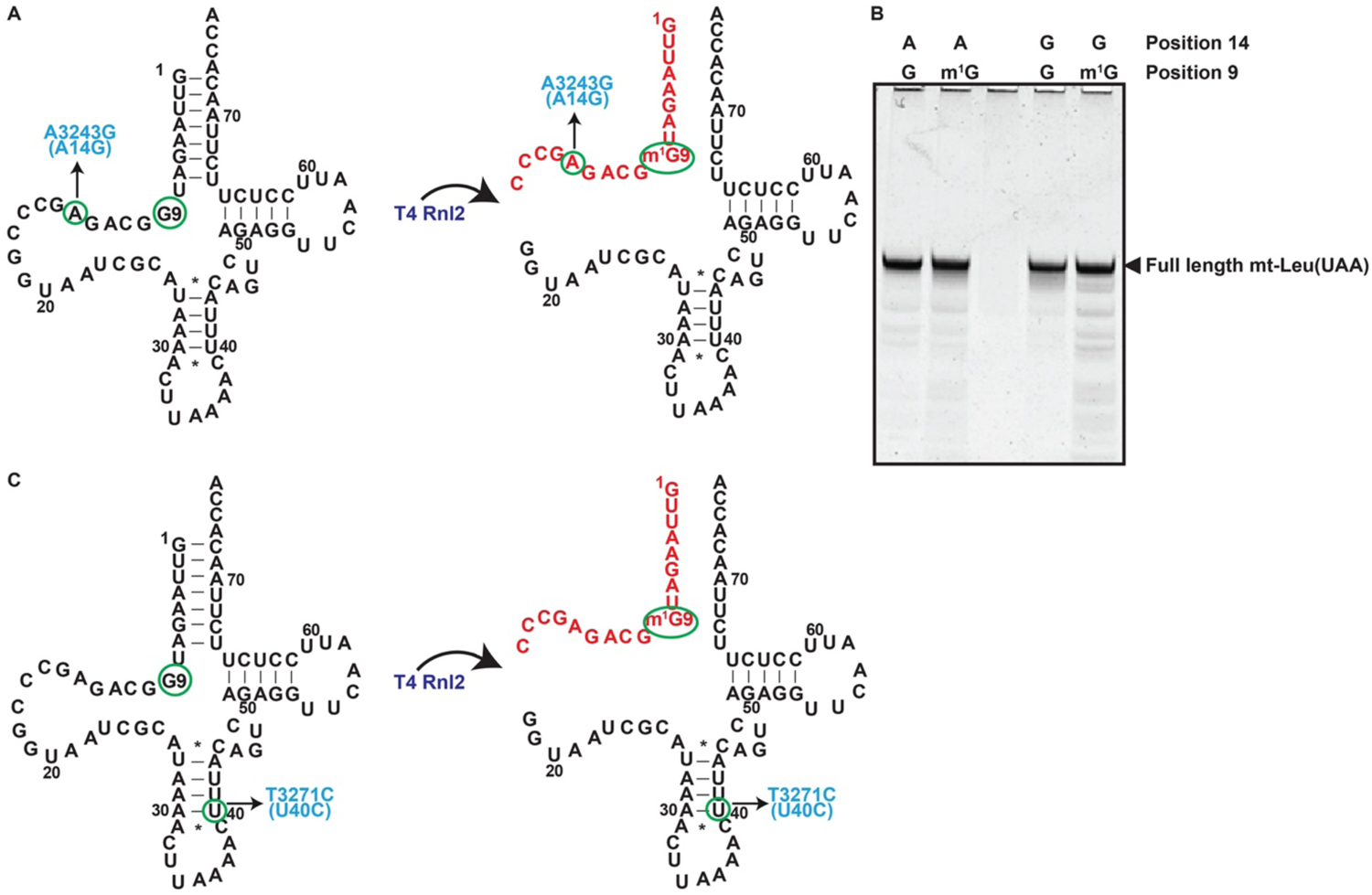
(Related to Figure 4) Assembly of mt-Leu(UAA) in the m^1^G9-state. (A) Synthesis of the G9-state of the transcript of WT and A3243G variant was by *in vitro* transcription, while synthesis of the m^1^G9-state was by ligation of the 5’-fragment (encoding nucleotides G1 to C17a) with the 3’-fragment (encoding nucleotides G18 to A76) using T4 Rnl2. The 5’-fragment was chemically synthesized, while the 3’-fragment was generated by *in vitro* transcription. (B) Denaturing gel analysis of the assembled full-length tRNAs of the WT and A3243G variant, in the G9 or m^1^G9-state of each. (C) Synthesis of the G9-state of the transcript of WT and T3271C variant was by *in vitro* transcription, while synthesis of the m^1^G9-state was by ligation of the 5’-fragment (encoding nucleotides G1 to C17a) with the 3’-fragment (encoding nucleotides G18 to A76) using T4 Rnl2. The 5’-fragment was chemically synthesized, while the 3’-fragment was generated by *in vitro* transcription.

**Figure S5.**
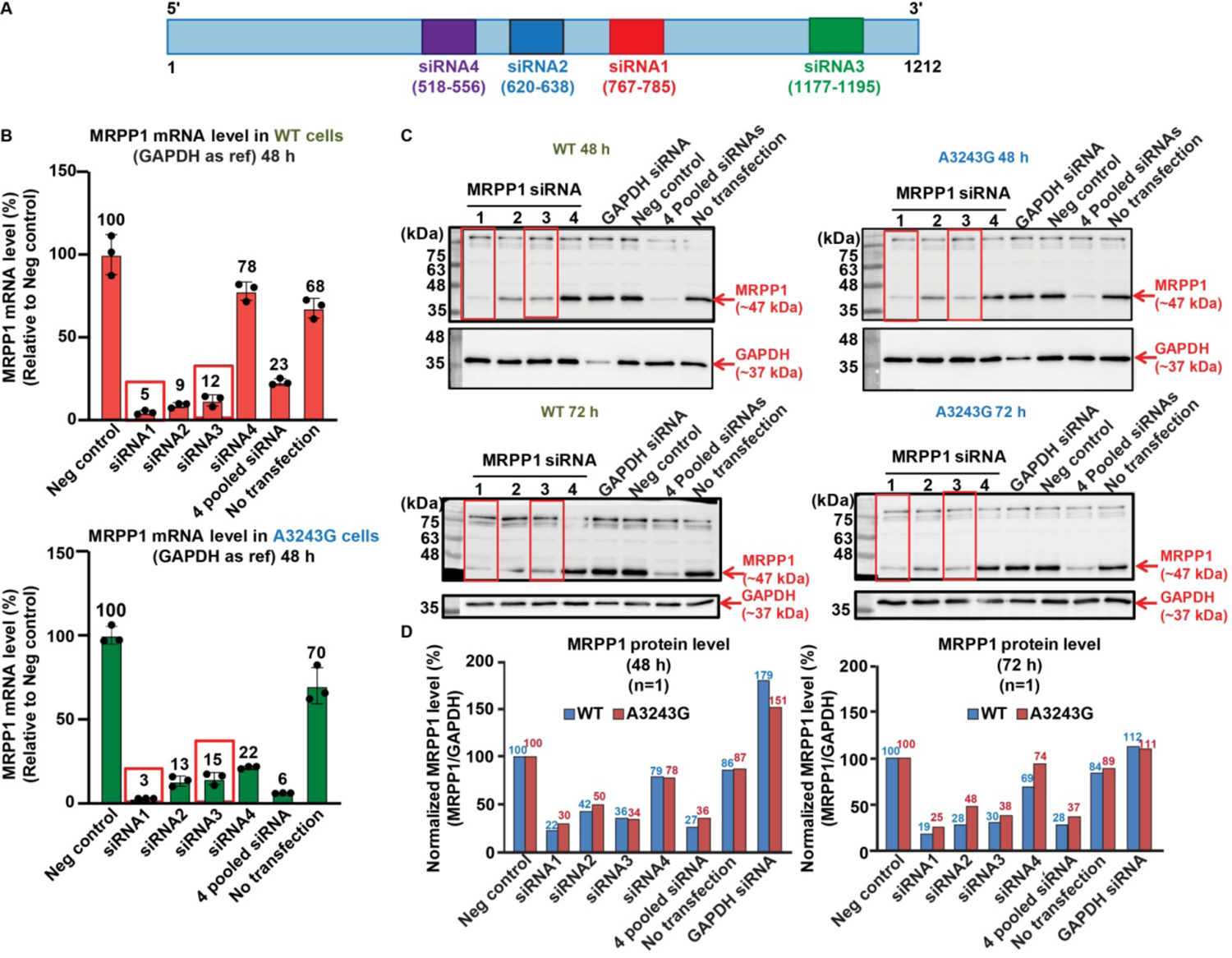
(Related to Figure 5) Targeting *MRPP1* mRNA by siRNAs (A) A linear map of *MRPP1* showing regions targeted by siRNAs. (B) Monitoring siRNA targeting of WT and A32434G cybrids at 48 h post-transfection by qRT-PCR analysis of the *MRPP1* mRNA level relative to the *GAPDH* mRNA. (C) Monitoring siRNA targeting at 48 h or 72 h post-transfection by Western blot analysis of the MRPP1 protein level relative to GAPDH. (D) Reduction of MRPP1 protein levels upon siRNA targeting at 48 h and 72 h post-transfection.

**Figure S6.**
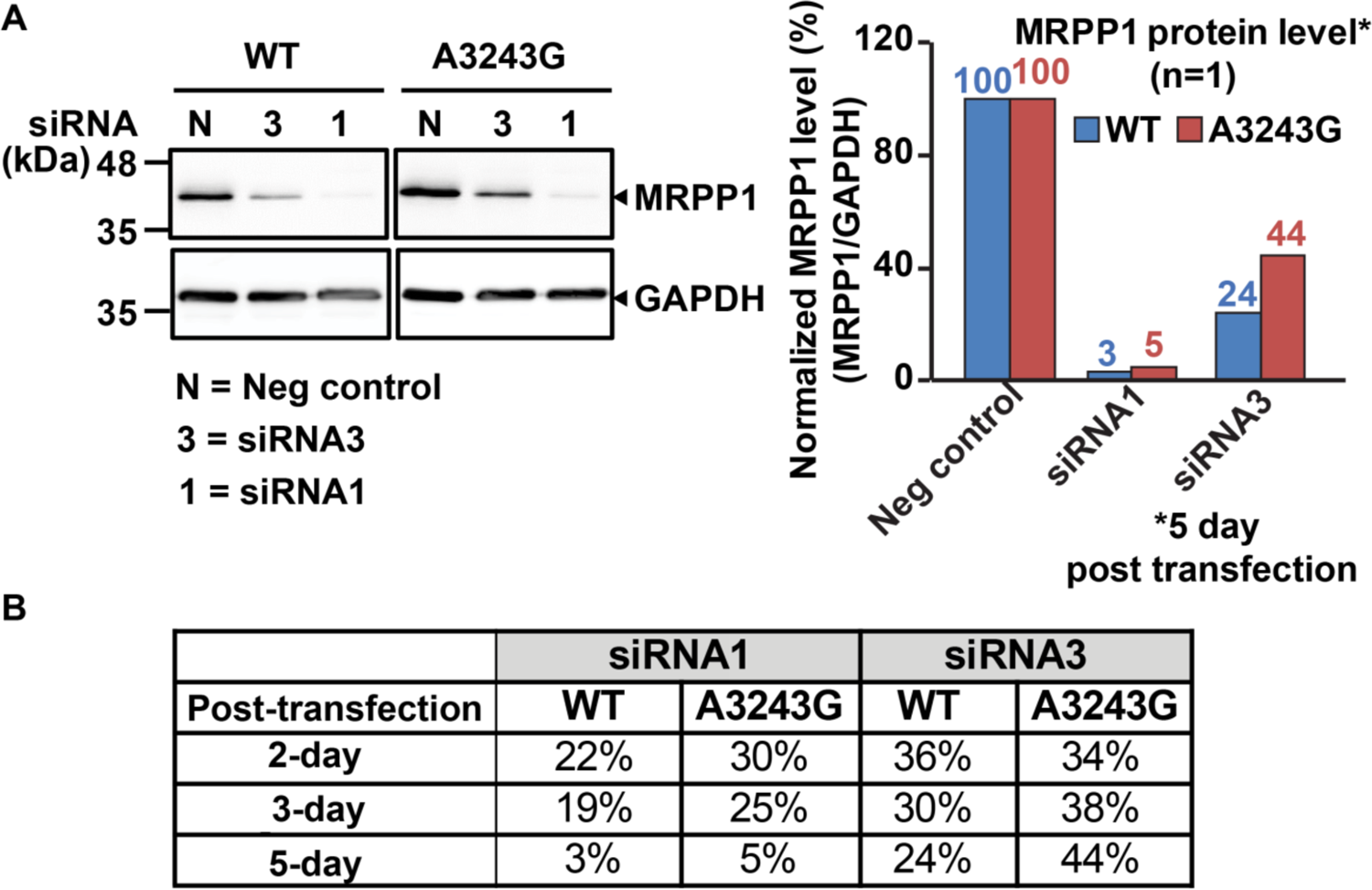
(Related to Figure 6) siRNA targeting of *MRPP1* in cybrids (A) The protein level of MRPP1 relative to GAPDH remained reduced 5-day post-transfection of cybrids with siRNA1 or siRNA3. (B) MRPP1 protein levels at 2-day, 3-day, and 5-day post-transfection of cybrids with siRNA1 or siRNA3.

## SUPPLEMENTAL TABLES

**Table S1.**
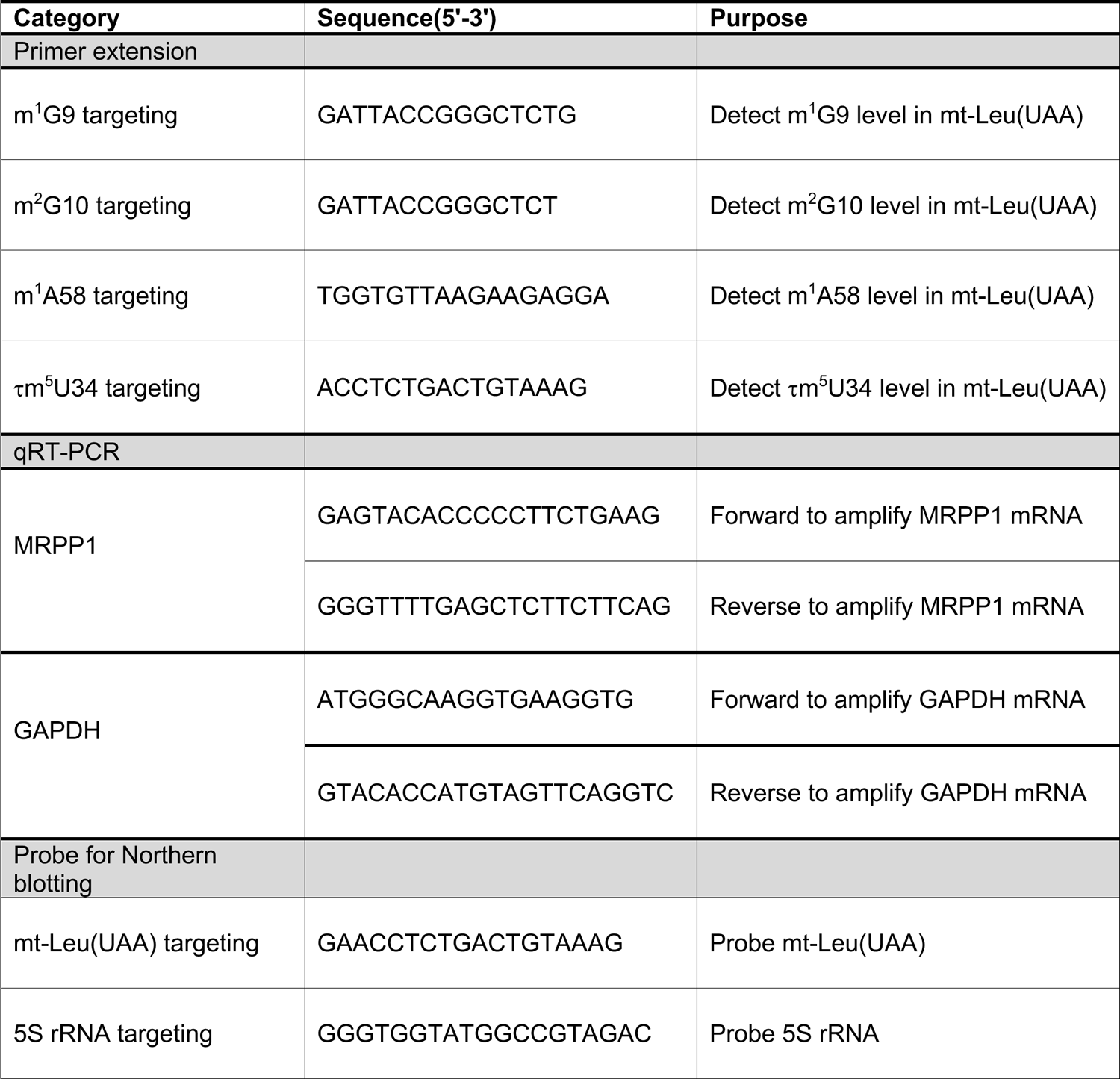
Primers used in this study.

**Table S2.**
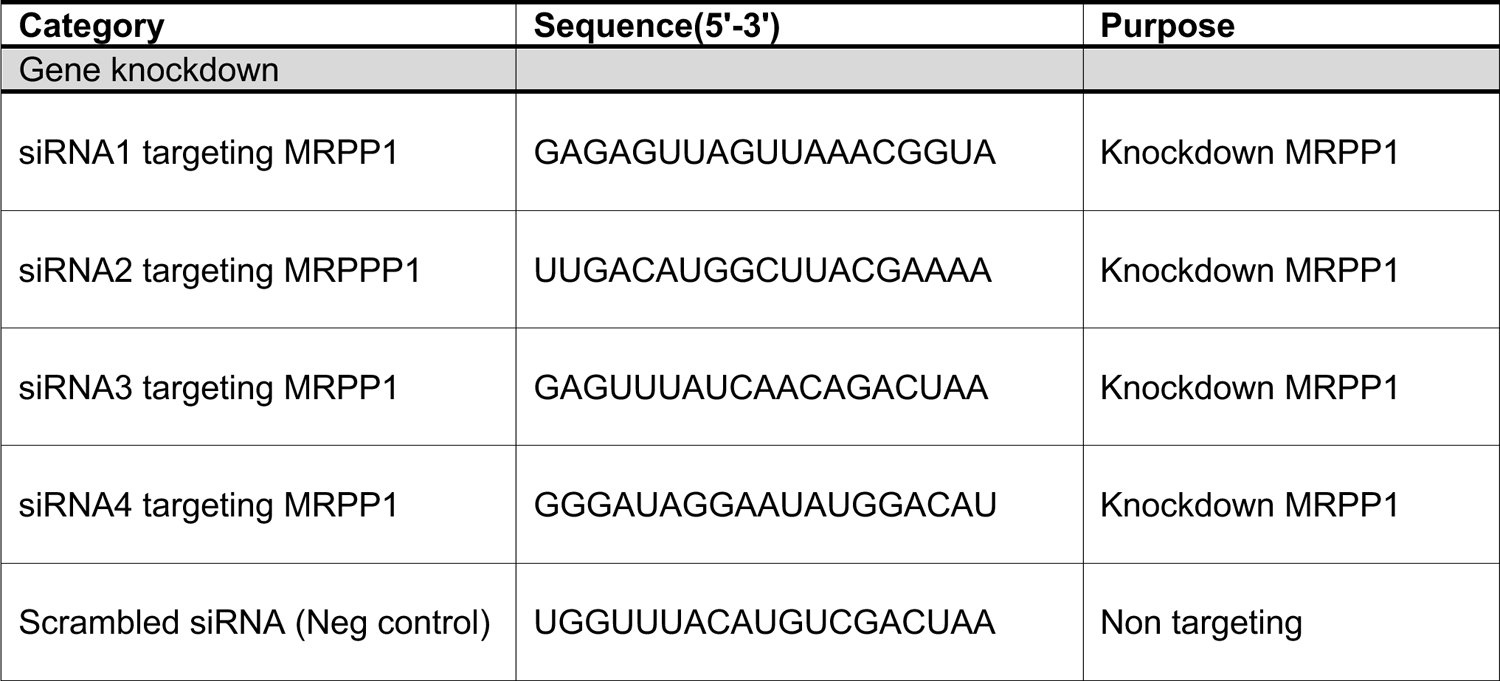
Targeting *MRPP1* by siRNAs.

**Table S3.** Quantification of tRNA stability (Related to **Figure 5D**, 5E)

|  |  | **Neg control** | **siRNA1** | **siRNA3** | **No transfection** |
| --- | --- | --- | --- | --- | --- |
| **mt-Leu(UAA)/5S rRNA** | **WT** | **0.44608** | **0.29067** | **0.33757** | **0.44516** |
|  | **A3243G** | **0.17009** | **0.14055** | **0.15122** | **0.16884** |
| **Values relative to WT Neg control (%)** | **WT** | **100** | **65.16** | **75.68** | **99.79** |
|  | **A3243G** | **38.13** | **31.51** | **33.90** | **37.85** |
| **A3243G values relative to its corresponding WT condition (%)** | **WT** | **100** | **100** | **100** | **100** |
|  | **A3243G** | **38.13** | **48.36** | **44.78** | **37.93** |

**Table S4.**
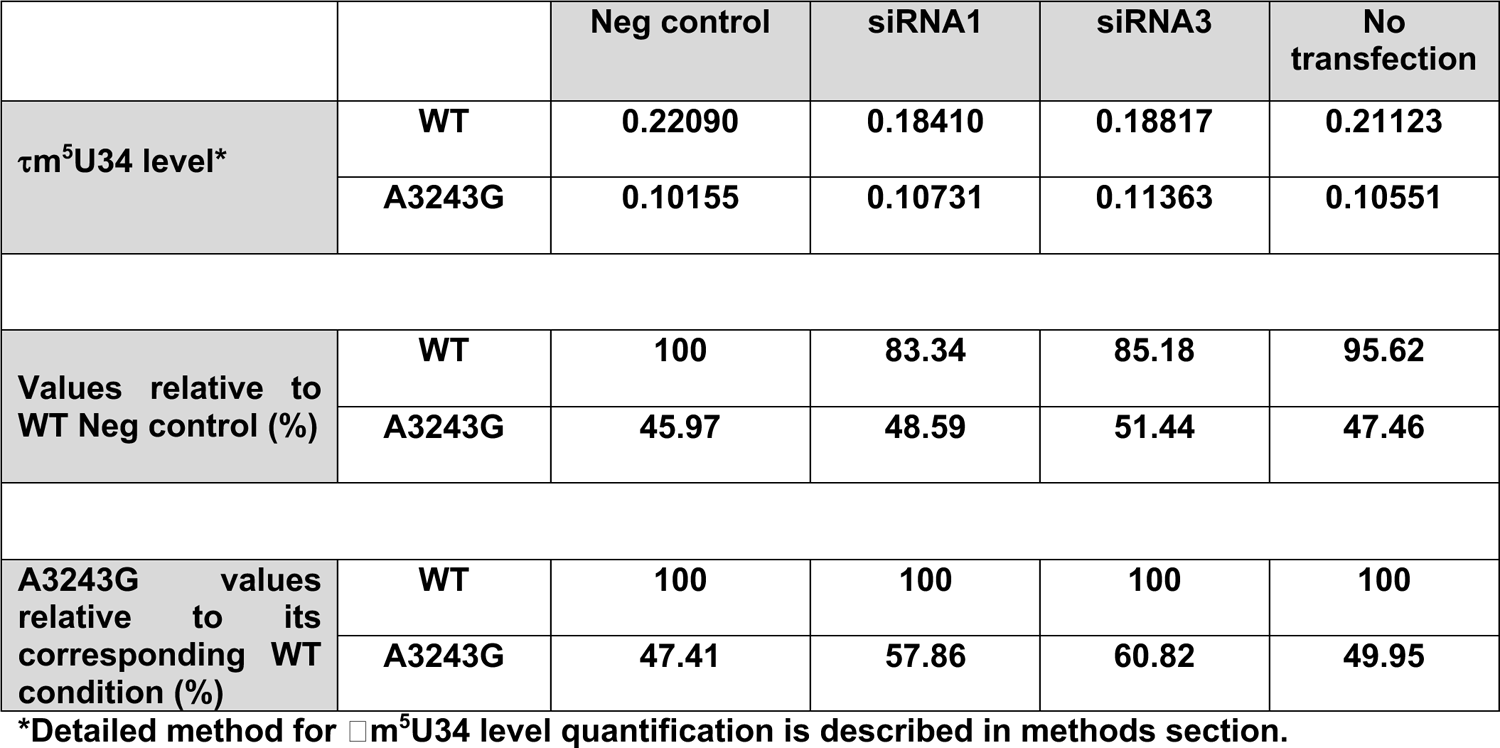
Quantification of 1m^5^U34 level (Related to **Figure 5G**, 5H)

**Table S5.** Quantification of charged tRNA level (Related to **Figure 5J**, 5K)f.

|  |  | **Neg control** | **siRNA1** | **siRNA3** | **No transfection** |
| --- | --- | --- | --- | --- | --- |
| **Charged tRNA level**<br>**(charged /**<br>**charged+Uncharged)** | **WT** | **0.83440** | **0.65314** | **0.64458** | **0.82566** |
|  | **A3243G** | **0.68924** | **0.70572** | **0.67035** | **0.68948** |
| **Values relative to**<br>**WT Neg control (%)** | **WT** | **100** | **78.28** | **77.25** | **98.95** |
|  | **A3243G** | **82.60** | **84.58** | **80.34** | **82.63** |
| **A3243G values**<br>**relative to its**<br>**corresponding WT**<br>**condition (%)** | **WT** | **100** | **100** | **100** | **100** |
|  | **A3243G** | **82.60** | **108.05** | **103.99** | **83.51** |

